# Assessing the Adequacy of Morphological Models used in Palaeobiology

**DOI:** 10.1101/2024.01.25.577179

**Authors:** Laura P. A. Mulvey, Michael R. May, Jeremy M. Brown, Sebastian Höhna, April M. Wright, Rachel C. M. Warnock

**Affiliations:** GeoZentrum Nordbayern, Department of Geography and Geosciences, Friedrich-Alexander Universität Erlangen-Nürnberg, Erlangen, Germany; Department of Evolution and Ecology, University of California Davis, Davis, CA USA; Department of Biological Sciences and Museum of Natural Science, Louisiana State University, Baton Rouge, LA, 70803, USA; GeoBio-Center, Ludwig-Maximilians-Universität München, 80333 Munich, Germany; Department of Earth and Environmental Sciences, Paleontology & Geobiology, Ludwig-Maximilians-Universität München, 80333 Munich, Germany; Department of Biological Sciences, Southeastern Louisiana University, Hammond, LA, 70402, USA

## Abstract

Reconstructing the evolutionary history of different groups of organisms provides insight into how life originated and diversified on Earth. Phylogenetic trees are commonly used to estimate this evolutionary history, providing a hypothesis of the events. Within Bayesian phylogenetics a major step in estimating a tree is in choosing an appropriate model of character evolution. In the case of most extinct species, our only source of information to decipher their phylogenetic relationships is through the morphology of fossils. We therefore use a model of morphological character evolution, the most common of which being the Mk Lewis model. While it is frequently used in palaeobiology, it is not known whether the simple Mk substitution model, or any extensions to it, provide a sufficiently good description of the process of morphological evolution. To determine whether or not the Mk model is appropriate for fossil data we used posterior predictive simulations, a model adequacy approach, to estimate absolute fit of the model to morphological data sets. We first investigate the impact that different versions of the Mk model have on key parameter estimates using tetrapod data sets. We show that choice of substitution model has an impact on both topology and branch lengths, highlighting the importance of model choice. Next, we use simulations to investigate the power of posterior predictive simulations for morphology. Having validated this approach we show that current variations of the Mk model are in fact performing adequately in capturing the evolutionary dynamics that generated our data. We do not find any preference for a particular model extension across multiple data sets, indicating that there is no ‘one size fits all’ when it comes to morphological data and that careful consideration should be given to choosing models of discrete character evolution. By using suitable models of character evolution, we can increase our confidence in our phylogenetic estimates, which should in turn allow us to gain more accurate insights into the evolutionary history of both extinct and extant taxa.

## 2 Introduction

The origination and subsequent diversification of species is a fascinating, yet complex, process. Phylogenetic trees serve as a powerful tool to aid in our understanding of this process. They provide a hypothesis of the evolutionary history of a group, enabling us to make inferences about the relationships, timing of events, and patterns of evolution (Baum and Offner, 2008). While molecular data may be more commonly used in phylogenetics (Lee and Palci, 2015), morphological data was the original source of evidence (Farris et al., 1970) and remains extremely valuable to our interpretation of species diversification (López-Antoñanzas et al., 2022). As the majority of life on Earth is now extinct, the fossil record contains a wealth of knowledge about how species have adapted and diversified through time (Simpson, 1952). Integrating this information into phylogenetic analysis, either in combination with molecular data, for example, in a total evidence approach (Gavryushkina et al., 2017; Mongiardino Koch et al., 2021) or independently, can therefore further our ability to resolve species relationships in deep time. Studies have also shown that incorporating fossil data into an analysis, even when the focus of the study is on extant taxa, can improve the topological resolution or even accuracy of a phylogenetic inference (Beck and Baillie, 2018; Koch and Parry, 2020; Mongiardino Koch et al., 2021). The use of morphological data in phylogenetics has been a topic of debate for many years, specifically, with regards to which approach should be applied, i.e., parsimony or model-based inference (Kolaczkowski and Thornton, 2004; Wright and Hillis, 2014; O’Reilly et al., 2016; Puttick et al., 2017; Sansom et al., 2018; Goloboff et al., 2018, 2019). Due to the complex nature of morphological data, there are doubts about our ability to correctly model its evolution, and that any assumptions made by the models will bias the resulting inference (Goloboff et al., 2019). Parsimony is often considered to be an assumption free approach; however, this is not entirely true, as there are still implicit assumptions about morphological evolution within a parsimony framework (Felsenstein, 1983; Steel and Penny, 2000). These two approaches have been compared many times throughout the literature, amassing in a large body of work which goes beyond the context of this study. Ultimately, model-based approaches have many more applications and statistical advantages, including the ability to select among competing models and assess model adequacy (Wright and Hillis, 2014; O’Reilly et al., 2016; Puttick et al., 2017). Amidst this debate, however, an important question has yet to be addressed: are available models of morphological evolution in fact adequate for our data?

Morphological data collected from fossils, or extant taxa, can be either discretized (e.g., presence/absence) or continuous (e.g., body size measurements). Discrete morphological data is the most widely used for phylogenetic inference (Lewis, 2001; Wright and Hillis, 2014; Harrison and Larsson, 2015; Wright, 2019) and will be the focus throughout this study. The data must be manually collected to create a morphological matrix, matching the format of a molecular alignment, where each site now represents a morphological trait. Traits are described using a character which is indicative of the phenotype expressed by a given taxon. Traits can have any number of character states depending on the complexity and traits with more than 2 states are referred to as multistate. Presence/absence traits can be described by using only 0 and 1, i.e., two character states. For more complex traits, however, more character states may be required. An example of this could be describing the shape of part of a skull or a shell. In this scenario a state is assigned to a particular modification of the trait, where a number of different adaptations (or states) may be present in a group. Within a single morphological matrix some traits can have binary character states, while others require multiple states. Consequently, the same character state across different traits can have an entirely different biological meaning, even within the same matrix. See Wright (2019) for a more in depth review of morphological data used in phylogenetics. The generation of this data is a challenging and time-intensive process, requiring an in-depth knowledge of the taxonomic group in question. Morphological data is, in turn, extremely valuable in helping us answer questions about the evolution of life that molecular data alone cannot answer.

Within a model-based phylogenetic analysis, the process that gives rise to discrete character data is described using a substitution model. These models aim to capture the evolutionary dynamics resulting in the gain, loss or modification of discrete states. Substitution models are continuous-time markov chain (CTMC) models. They allow states to change (evolve) stochastically at any point in time, and this change depends only on the current state that the evolving system is in. The assumptions of a substitution model are mathematically represented using a Q(or rate)-matrix. A Q-matrix is a square matrix where each element represents the instantaneous rate of change between states. That is Q[*i, j*] represents the rate of change from state *i* to state *j*. The probability of change over a given interval, or branch length *v*, is calculated using the Q-matrix. Developing models that can accurately describe the complex processes driving morphological evolution is extremely challenging and as a result, there is only one main model that is commonly applied: the Mk model (Felsenstein, 1992; Lewis, 2001). This model is a generalisation of the Jukes Cantor model (Jukes and Cantor, 1969) used for molecular data, and as such, follows the same set of assumptions. It assumes equal transitions rates between states, that is, the probability of transitioning from a state 0 to a 1 is the same as going from a state 1 to a 0. It also assumes equal base (state) frequencies, meaning the model expects that there is approximately the same number of each character state throughout the morphological matrix. The Q-matrix for such a model, therefore sets all transitions to have an equal probability, with its size being determined by the number of states. That is, for a purely binary data set the Q-matrix will be a 2×2 matrix, representing the transitions from state 0 to state 1, from state 1 to state 0, and of no change.

Morphological data is, needless to say, different to molecular, so there are concerns about how well a model originally developed for molecular data can be applied to morphological data. Additionally, given that more complex models are often selected for molecular data, there is doubt about how well such a simple model can be applied to morphological data. As such, there have been a number of extensions implemented for the Mk model to relax these strict assumptions, and allow the model to better describe the reality of morphological evolution. Lewis immediately noted an important difference between morphological and molecular data collection (Lewis, 2001). When taxonomists are creating a matrix, or character coding, they will typically exclusively choose traits which differ across species, resulting in a matrix where every site is variable. This is a markedly different behavior from molecular data collection, where there can be many sites where a nucleotide is conserved across all species. Not accounting for this phenomenon, known as ascertainment bias, (though referred to as acquisition bias in Lewis (2001)), can result in inferring trees with extremely long branch lengths. Lewis dealt with this by conditioning the likelihood calculation on there only being variable characters, developing the MkV model. There are a number of other extensions that we will explore the effects of here as well. Accounting for among-character rate variation has also been suggested as important when modeling morphological evolution (Harrison and Larsson, 2015). This allows different traits to transition at different rates, as some may be evolving faster than others. This is frequently achieved by drawing rates from a discretized gamma distribution and allowing a trait to transition according to a given rate category, the same as is done for molecular data (Yang, 1994). Data sets can also be partitioned, often based on the maximum number character states (e.g., see Khakurel et al., in press). This ensures that traits are in a Q-matrix of the correct size. That is, in an unpartitioned analysis, the Q-matrix will take the size of the maximum character state in the morphological matrix, which could be for example 5. Transitions between binary characters will therefore also be calculated in this Q-matrix of size 5, meaning that there is some probability given to a binary character of transitioning to states 2, 3, or 4. As we do not observe these states in the data, in some cases (e.g., where states 1 or 0 are used to represent presence or absence) we can be certain that this is incorrect. Partitioning by character states such that all binary characters are in a Q-matrix of size 2 and so on, avoids this issue. Partitioning data can have an effect on branch lengths (Khakurel et al., in press) so it is important that it is done when necessary. Similarly, however, incorrect partitioning may lead to too low rates as a result of observer bias.

The impact of these different variants of the Mk model is still not fully understood in terms of the effects on key parameter estimates, although they are likely to cause differences as has been shown for molecular models (Lemmon and Moriarty, 2004). When deciding what model to use, there are two distinct questions that can be asked, (1) which is the best model for my data compared to other models? and/or (2) does this model fit my data? The first question, which is the more common of the two, can be answered using model selection. Model selection approaches are common in molecular based studies although less frequently used for morphological data. For morphological studies there is a history of using substitution models that have been used in previous studies, choosing a model based on the structure of the data set, or relying on software defaults, often without providing statistical justification for model choice. As previously stated, data sets are manually produced, meaning they can differ from each other depending on the taxonomist. If, for example, a substitution model had been applied to the taxonomic group of interest in the past, even if you are using similar taxa, if the morphological matrix is different, using the same substitution model as previous studies may not be logical. That being said, there are a number of examples where model selection has been applied to morphological data sets (Caldwell et al., 2021; Rücklin et al., 2021; Wright et al., 2021). By using a model selection approach, any subjectivity in model choice can be reduced. One down side of model selection approaches, however, is that they give no indication of the absolute fit of the model to the data. It tells you which model is the relative best, but that does not necessarily mean that the model provides a good description of the true data generating process, simply it fits better than other models (Gatesy, 2007). This is where question two becomes important. Asking if a single model is adequate allows you to understand how well a model can describe your data. These approaches, known as model adequacy, are currently gaining in popularity for molecular data (Duchêne et al., 2017, 2018; Brown and Thomson, 2018) and have been sporadically applied to morphological data sets (Huelsenbeck et al., 2003; Slater and Pennell, 2014) but have yet to be systematically assessed.

In order to confidently integrate fossils into phylogenetic approaches, ensuring we have accurate substitution models is a critical step. Knowing that the models are behaving as expected can increase our confidence in the results and allow us to ask increasingly complex questions. Here we explored the impacts of different substitution models on key parameter estimates across a number of morphological data sets, as well as investigated the best approaches for choosing a model. We found that the models have a notable impact on both tree length and topology, highlighting the importance of validating a model before using it. In our simulation study, model adequacy preformed well in predicting which model the data was simulated under. Ultimately, using model adequacy, we found that substitution models do in fact fit a number of empirical data sets, supporting the use of the Mk model for morphological data in paleobiology.

## 3 Methods

### 3.1 Data

We used a collection of previously published morphological matrices from Sansom et al. (2018) (taken from http://graemetlloyd.com/matrdino.html). This data set contained 166 morphological matrices of tetrapod taxa. The data sets vary in sizes in terms of taxa, from 12-219, traits, from 23-622, and number of different character states, from 2-10. They have also been used previously to examine the use of phylogenetic methods and as such were a ideal data set for this study Sansom et al. (2018). We removed matrices based on two criteria: (i) those that contained characters with more than 9 states or 80 taxa, as they became too computationally expensive, and (ii) those that contained traits where only character state “0” and missing characters, “?” were present for any trait. This resulted in a final data set of 114 matrices. The data sets varied in size, with the number of taxa ranging from 12 to 80, and the number of characters being between 23 to 477.

### 3.2 Empirical Comparison of Morphological Models

Initially, our focus was on investigating how substitution models impact the estimation of key parameters. We chose 7 variants of the Mk model (Mk, MkV, MkV+G, Mk+G, MkVP, MkVP+G, MkP+G, see Table 1 for model assumptions) and compared differences in the resulting tree lengths and topologies. All phylogenetic inference was performed in a Bayesian framework using the software RevBayes version (1.2.1) (Höhna et al., 2016). We ran an MCMC inference under each of the 7 models for all 114 data sets. This allowed us to determine whether there are any systematic differences in parameter estimates that could be attributed to the substitution model. For all models we assumed a uniform tree prior on the topology. Tree length was drawn from an exponential prior distribution with a rate parameter of 1. Relative branch lengths were drawn from a Dirichlet prior distribution (Zhang et al., 2012). The branch lengths were calculated as the product of the tree length and the relative branch lengths. Preliminary analyses were run using an exponential prior for branch length estimation, however, we found the Dirichlet tree prior to perform better in simulations. We used an Mk model, with the size of the Q-matrix being determined by the maximum character state of each data set. When allowing for among character rate variation, ACRV, (+G) the shape parameter of the gamma distribution, *α* was estimated as the inverse of a random variable alpha inv drawn from the exponential distribution with a rate parameter of 1. We discretized the gamma distribution into four discrete categories (Yang, 1994). To account for ascertainment bias (+V), we selected the variable coding option in RevBayes. Partitioned models (+P) split the data set based on the number of character states. Each grouping had its own Q matrix. That is, all binary traits were assigned to a Q-matrix of size 2, all tertiary traits were assigned to a Q-matrix of size 3 and so on. For this set up, we applied the same gamma distribution for ACRV to each partition.

**Table 1:** Models tested.

| Models & Extensions | Assumptions |
| --- | --- |
| Mk | all transition are equal (Lewis, 2001) |
| V | accounts for ascertainment bias (Lewis, 2001) |
| G | allows for variation in substitution rates among sites (Yang, 1994) |
| P | partitions the data based on the number of character states |

We ran the MCMC for 20,000 iterations with two simultaneous chains, sampling every 10 generations. The output of both chains was automatically combined in RevBayes, resulting in a posterior sample of 4,000. Convergence was assessed using a custom R script with the R package coda (Plummer et al., 2006) to ensure ESS values *>* 200 of all parameters estimated.

#### 3.2.1 Posterior Summaries

Tree length was calculated as the sum of the branch lengths averaged across the entire posterior distribution. We also calculated the percentage change in tree length relative to the Mk model for each data set to make it easier to observe any consistent patterns across models. We then explored the differences in estimated tree topologies from the different substitution models for each data set. Using a sample of 1000 trees from the posterior distribution for each substitution model, we calculated the normalised Robinson-Foulds distance between all trees. With this resulting matrix we performed a multivariate homogeneity of group dispersions analysis using the R package vegan (Oksanen et al., 2022). This calculated the distance between points and their group centroid. Plotting this as a PCoA allowed us to visualise where models were in tree space, relative to one another. In order to quantify these differences, we carried out a permutation test to assess their significance using the permutest function in the vegan package (Oksanen et al., 2022). This allowed us to determine if the variability in RF distances inferred using each of the models was significantly different from each other.

### 3.3 Assessing the Performance of Model Adequacy and Model Selection Methods for Morphological Data

Choosing an appropriate model of evolution is an important step in any Bayesian phylogenetic analysis. The results from an inference will be conditioned on the assumptions of the evolutionary model. As such, if the model’s assumptions are markedly different than that of the underlying process that generated the data, the results may be inaccurate. Methods for choosing an appropriate model often take a model selection approach, relying on estimation of the marginal likelihood (Brown, 2014b). These methods provide the relative fit of competing models. Although a model may be selected as the best choice, it does not necessarily mean that the model is, in any way adequate for the data set being analysed. That is, it may not provide a sufficiently realistic description of the data generating process (Gatesy, 2007; A Shepherd and Klaere, 2019). Therefore, model selection provides no indication about how well the model actually fits your data, only its relative fit compared to other models. In contrast, model adequacy approaches provide information on the absolute fit of a model to a data set. They can provide information about a model’s ability to capture key characteristics of a given data set, as well as highlight where the model may be inadequate. Importantly, model adequacy provides the ability to reject models, even if they are identified as the “best” using a model selection approach (A Shepherd and Klaere, 2019; Brown and Thomson, 2018).

Posterior-predictive simulations (PPS) is a model-adequacy approach that has been applied to a variety of data types, albeit with limited frequency in phylogenetics (Gelman et al., 1996; Bollback, 2002; Brown, 2014a; Brown and Thomson, 2018; Höhna et al., 2018; Schwery et al., 2023). Briefly, it works by simulating data under a given model and comparing the similarity of the empirical data to the newly simulated data using a test statistic. The rationale here being that if the model adequately captures the underlying dynamics of the processes generating the data, the simulated data would be similar to the empirical (Gelman et al., 1996; Bollback, 2002). To date, the use of PPS has been demonstrated more often for molecular data, for example Brown (2014a) and Duchêne et al. (2018), however, it has also been suggested for models of continuous trait evolution (Slater and Pennell, 2014), and discrete character evolution (Huelsenbeck et al., 2003). Using simulations, we investigate the use of Bayes factors and PPS for determining whether a morphological model fits our data.

#### 3.3.1 Model Adequacy Using Posterior Predictive Simulations

To test the adequacy of morphological models we used posterior prediction simulations (PPS) following the workflow as described in Höhna et al. (2018) implemented in RevBayes. This can be broadly broken down into four mains steps. We provide a brief description of these steps here, but for a more thorough description see Höhna et al. (2018). (1) The first step is to analyse the empirical data under a given model. This involves a regular MCMC inference sampling parameter values from the posterior distribution. (2) New data sets are then simulated in R using the phangorn R package (Schliep, 2011). Data sets are simulated under the same model as used in step 1 with trees and parameter estimates inferred in step 1. (3) Inference under the same model is then carried out on all the newly simulated data sets from step 2. (4) Test statistics are calculated and compared between the original empirical data and inference results, and the newly simulated data and inference results, see Fig. 1. The overarching idea here being, the more similar the simulated data is to the empirical data, the better the model is at describing the underlying processes that produced your data. This in turn indicates whether we can have confidence in the results inferred under a given model. Note it is practical to simulate data sets in RevBayes, and we provide instructions for doing so in the associated tutorial (https://revbayes.github.io/tutorials/pps_morpho/). We chose to simulate data using phangorn as it was slightly more computationally efficient given that our study featured an exceptionally large number of simulations (700,000 simulations for 160 individual data sets), but this should not be a concern for an empirical study, which would typically only contain one or a few individual data sets.

**Figure 1:**
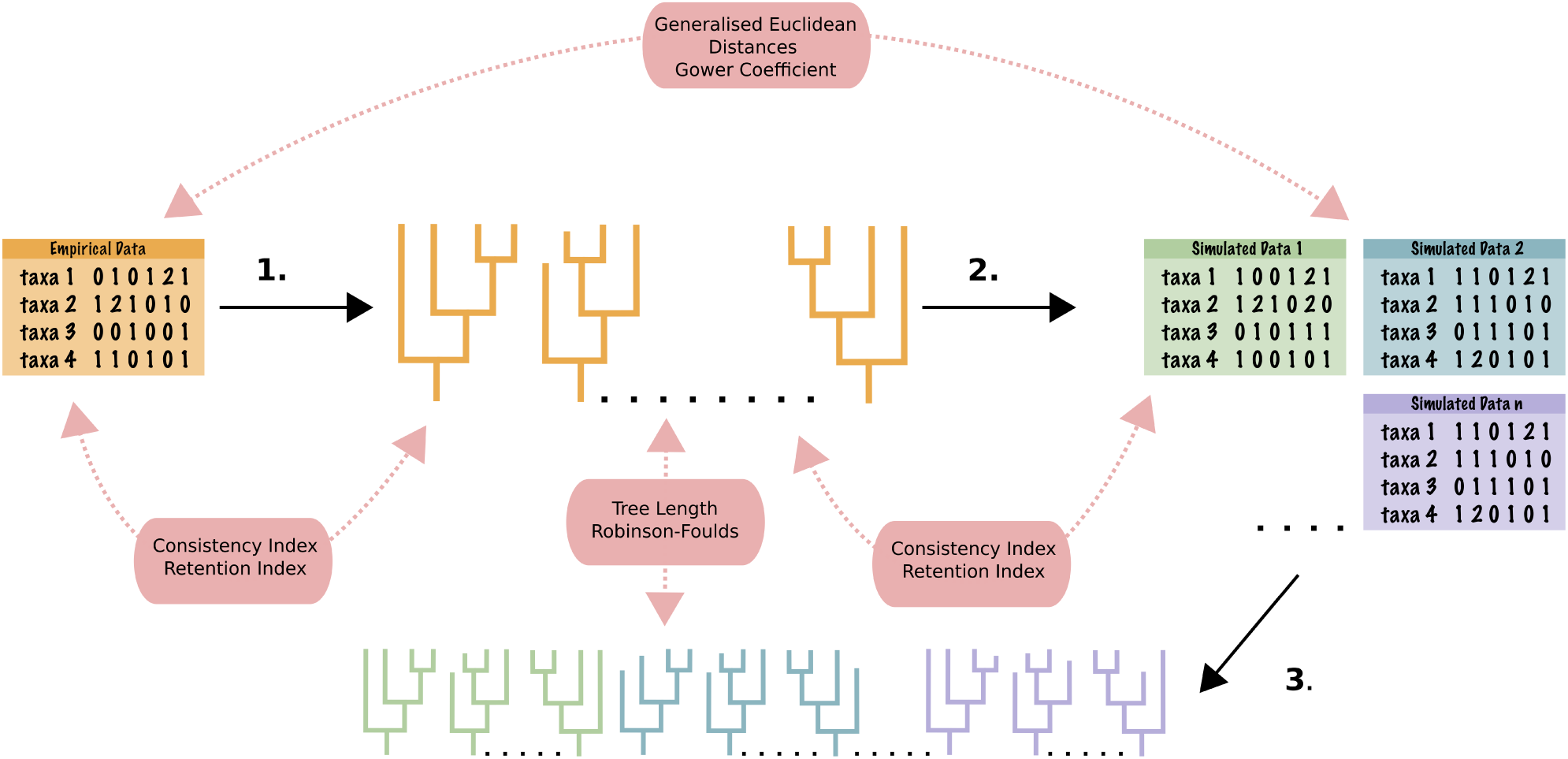
Posterior predictive simulation workflow. **Step 1.** an MCMC inference is carried out under a given model. **Step 2.** data sets are simulated under the same model based on parameter estimates from 1. **Step 3.** an MCMC inference is then carried out on the simulated data sets. The pink boxes show the test statistics that are applied to determine whether or not the model is adequate. Generalised Euclidean distances and Gower’s coefficient are used to compare the data sets. Tree length and Robinson-Foulds are used to compare the inferred trees. Consistency index and retention index use the empirical trees and the empirical and simulated data sets to test for adequacy.

#### 3.3.2 Candidate Test Statistics for Morphological Data

PPS are only as good as the test statistics used, meaning if the test statics are not able to capture differences that result from the underlying dynamics of the data generating processes, it will not be possible to use PPS to understand the adequacy of a given model. Using test statistics allows us to convert the empirical data and output into numerical values that we can use to summarize the differences between empirical and simulated data. The test statistics can then be compared using effect sizes, which provide a way of quantifying variation in model fit and allow us to distinguish between the fit of competing models. Previous studies have used posterior-predictive *p*-values to accept or reject a model. In this study we chose to focus on effect sizes over *p*-values for two reasons. First, given that fit of morphological models to empirical data had not been tested previously, we wanted to determine how different models preformed and essentially, potentially how poorly they each fit empirical data. Second, effect sizes provide a more intuitive way of comparing the fit of different models. By applying *p*-values only we can assess whether a model is adequate or not, but not how the models perform relative to each other (Brown, 2014a; Duchêne et al., 2017). Effect sizes therefore allow us to gain a better understanding of the impact of different morphological models, and ultimately address the main questions of this study. This would not be necessary perhaps in an empirical study, and we do include the use of *p*-values for our empirical analysis.

Here, the effect sizes were calculated by:

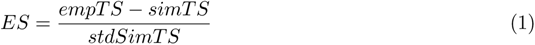

where *empTS* is the empirical value for a given test statistic, *simTS* is the value of the test statistic from a single simulated replicate and *stdSimTS* is the standard deviation across all simulated replicates. The closer this number is to zero, the better the model is at explaining your data. Test statistics can be divided into three categories: (1) data based, (2) inference based, and (3) data inference hybrid or mixed. Data based test statistics compare the actual morphological data sets themselves, inference based compare the inferred trees and mixed statistics uses both the data and the trees to compare your empirical and simulated values.

##### Data Based Test Statistics

As the name suggests, these test statistics focus on characterising the matrices, themselves here meaning the morphological data. As PPS studies in phylogenetics have previously focused on molecular data, many of the data based statistics are only suited to DNA. For example, quantifying the GC content or number of invariant sites (Höhna et al., 2018). Summarising morphological data sets in a similar way requires different metrics. To do this we explore the use of disparity metrics. Disparity is a measure of the morphological variation observed among species (Hopkins et al., 2017). It is important to note, we are not interested in the actual measure of disparity, we are interested in how the value differs between the original empirical data and the simulated data. We tested two metrics of disparity.

(i) *Generalised Euclidean Distances* (GED) (Wills, 1998) is a popular disparity metric commonly used in vertebrate research (Brusatte et al., 2011; Lehmann et al., 2019). This measure is similar to the basic Euclidean distances but incorporates adjustments to accommodate missing characters. Wills (2001) defines GED as:

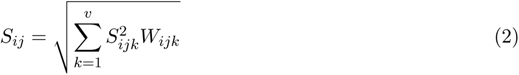

where *S _ij_* is the total distance between taxa *i* and *j*, *v* is the total number of characters in the matrix, *W _ijk_* is the weight of the *k* th character, and *S _ijk_* is the distance between taxa *i* and *j* at the *k* th character. *S _ijk_* equals 0 when the *i* th and *j* th sequence match in the *k* th position and 1 when there is a mismatch. To account for missing data, a mean estimate of disparity is first calculated across all comparisons for which we have observations:

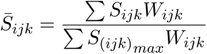

where *S*_(_*_ijk_*_)_ is the maximum possible distance between taxa *i* and *j* for the *k*th character, which equals 1 for discrete characters. The term 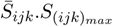 is then substituted into Equation 2 for missing *S_ijk_* values. In all cases, we treat characters as equally weighted, i.e., *W_ijk_* = 1.

(ii) *Gower’s Coefficient* (GC) (Gower, 1971) is commonly used in invertebrate studies (Hopkins and Smith, 2015). This metric calculates disparity differently to the GED, notably in regards to how it deals with missing characters. Here this is achieved by normalising by the available data. GC can be written as (Lloyd, 2016):

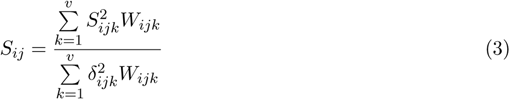

where *δ_ijk_* is coded as 1 if both taxa *i* and *j* can be coded for *k* (i.e., character states are observed for both taxa), and zero if not. As above, we use assume equal weights, i.e., *W_ijk_* = 1.

For both the above metrics, we used the R package Claddis (Lloyd, 2016). In the calculations we set characters as unorderd. The output from this matrix of the pairwise distance between taxa. We took the average disparity across the matrix for the calculation of the effect size, i.e., for *empTS* and *simTS*.

##### Inference Based Statistics

Inference based test statistics aim to characterise the inferred trees in the posterior distribution.

(i) *Mean Tree Length* (TL) was calculated using all the tree lengths sampled in the posterior distribution as:

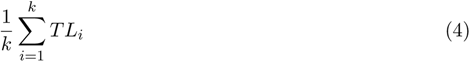

where TL is defined as the sum of branch lengths 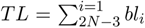. This calculation was done in RevBayes. We took the mean tree lengths across the posterior distribution of trees as the input for the effect sizes.

(ii) *Mean Robinson-Foulds Distance* (RF) was used to measure the topological uncertainty within the posterior distribution (Robinson and Foulds, 1981). This value was calculated in RevBayes.

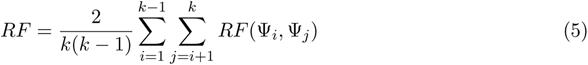

##### Mixed Test Statistics

These test statistics take both the data and the tree into consideration. Again, we investigate the use of two test statistics here.

i. *Consistency Index* (CI) (Kluge and Farris, 1969) which is a measure of homoplasy within the data set. It can be calculated as:

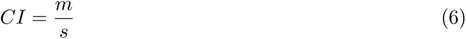

where *m* is the minimum possible number of steps or changes along a tree and *s* is the reconstructed number, i.e., the number observed along estimated trees (Kluge and Farris, 1969). This metric has been used to characterise data sets in paleontology (Murphy et al., 2021) and has been applied to model adequacy studies focusing on molecular data (Duchêne et al., 2018). A CI of 1 indicates no homoplasy and gets closer to zero as the amount of homoplasy increases.

(ii) The *Retention Index* (RI) (Farris, 1989), builds on the consistency index to calculate the potential synapomorphy observed along the tree and is calculated as:

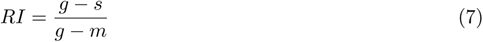

where *g* is the maximum number of possible steps on a given tree.

For both consistency and retention index, we used the maximum clade credibility (MCC) tree generated from inference of the empirical data for all calculations. We carried out preliminary analysis where we used the entire posterior distribution of trees for this calculation. The increased computation time from a number of minutes to 24 hours and produced extremely similar results, see fig. S2. For this reason, we continued to use the MCC tree only for the rest of the analysis.

#### 3.3.3 Model Selection Using Stepping Stone Sampling

For model selection, Bayes factors are computed to compare between models. In order to do this we first have to calculate the marginal likelihood of the data. The marginal likelihood is an important quantity in Bayesian model selection as it provides a measure of the goodness of fit of the model to the data, while accounting for model complexity. The marginal probability is the probability of the data integrated over all possible parameter values weighted by their prior probabilities for a given model. This is tricky to calculate so we avoid calculating it in regular MCMC inference using the Metropolis-Hastings algorithm (Metropolis et al., 1953; Hastings, 1970). We therefore, need to use a different approach in order to approximate this value. One such approach is stepping stone sampling. Stepping stone sampling is a Monte Carlo method that uses a sequence of intermediate distributions, or steps, between the prior and posterior distributions to compute the marginal likelihood. Stepping stone sampling has been demonstrated to be a reliable method for calculating Bayes factors and therefore performing model selection with molecular data (Xie et al., 2011; Höhna et al., 2021). While comparing marginal likelihoods has been used for morphological data to choose a model, its performance has yet to be assessed (Wright et al., 2021).

#### 3.3.4 Simulated Data

We based our simulation study on two empirical data sets, one on Proboscideans (the group containing elephants and their nearest extinct relatives) (Shoshani et al., 2006) and the other on Hyaenodontidae (Egi et al., 2005). For simplicity we will refer to each data set as simulated elephants and simulated hyenas, respectively. The simulated elephant data set is larger, having 40 taxa, 125 characters with 6 states compared to the simulated hyaenas which has 15 taxa, 65 characters and 5 states. For each data set, we used 20 trees from the posterior distribution inferred under a given model and simulated character data under the same model in R using phagnorm Schliep (2011). We did not simulate any traits with missing data. We did this for the MkV, MkVP, MkV+G and MkVP+G models for each data set (160 simulated replicates in total).

#### 3.3.5 Analysis of Simulated Data

We carried out PPS following section 3.3.1 on all simulated elephant and simulated hyena data sets. This allowed us to jointly validate the candidate test statistics and determine how well PPS can detect the correct model, as well as how it handles incorrect models. We analysed each of the simulated data sets under the same seven models as in section 3.2 (Mk, MkV, MkV+G, Mk+G, MkVP, MkVP+G, MkP+G) and kept all model parameters the same. The MCMC was ran for 10,000 iterations, with two individual chains. Convergence was assessed by calculating the ESS values for the likelihood, prior, posterior, tree length and when present in the model, the estimated alpha values using the R package coda (Plummer et al., 2006). MCMC chains that produced ESS values *<* 200 were ran again with an increase in the chain length. There were 560 replicates for each data set size. For the simulated hyena data sets, 533 converged after 10,000 iterations, 24 after 50,000 iterations and 3 after 100,000. For the simulated elephant data, 548 reached convergence after 10,000 iterations and 12 required 50,000 iterations.

The number of simulations required for PPS is not strictly defined. Given that the number of simulation replicates will increase both the computation time and memory requirements, doing extra should be avoided. To explore this we used both of the simulated data sets, simulated under the MkV+G model. We ran an MCMC inference as described above with 1,000 simulation replicates. We calculated the cumulative means for each test statistic inferred under each model. Following Robinson et al. (2004), we plotted the cumulative means thereby taking a graphical approach that shows the point at which the line becomes flat, indicating the required number of replicates Fig. S3, (Robinson et al., 2004). We found that after 500 replicates the lines were flat and we determined this to be sufficient. To ensure that this number of simulation replicates was not effecting the calculation of the actual effect sizes, we compared the effect sizes for each test statistic with 500 and 1,000 replicates. For ~92% of the effect sizes calculated, we found that the difference was less than 0.1 with a median of ~0.03. The largest change in effect sizes we saw was between 500 and 1,000 replicates which was 0.5. This was calculated for the two data based test statistics both inferred under the model MkVP+G and the same replicate. This results was thus considered an outlier. All other differences were less than 0.25, and did not change whether a model was considered to be adequate or not. As a results of these tests, we determined that having 500 simulations replicates would be sufficient for our PPS analyses throughout.

We then used stepping stone sampling to estimate the marginal likelihoods under each of the models. We kept all model parameters the same as above, and used 48 stones.

### 3.4 Analysis of Empirical Data

Once we identified appropriate test statistics, we could test model fit using PPS on empirical data sets to determine which, if any, morphological models were adequate. We chose to analyse 8 data sets here. This was limited by the computational costs of running the analysis multiple times. Data sets were chosen to cover a range of sizes, in terms of taxa, characters and states. We tested the same 7 models we used throughout (Mk, MkV, MkV+G, Mk+G, MkVP, MkVP+G, MkP+G) and kept all model parameters the same as in section 3.3.1. We also used stepping stone sampling on each of the data sets in order to see how the models chosen by model selection compared to those identified as most appropriate by model adequacy. Posterior *p*-values were calculated in R for each of the test statistics to compare with the results obtained using effect sizes.

## 4 Results

### 4.1 Empirical Comparison of Morphological Models

Assuming different models of morphological evolution produced different estimates of key parameters of interest. Figure 2A shows the percentage difference in mean tree lengths relative to that of the Mk model for all 114 data sets. There are some general trends that emerged here. As expected (Lewis, 2001), the MkV model produced smaller estimates of tree length relative to the Mk model for all but one data set. The Mk+G model produced longer trees for 96% of the data sets compared to the Mk model. However, when used in combination, these two extensions produced the smallest trees compared to all models in 96% of data sets. Partitioned models estimated larger trees, with the MkP+G model estimating larger trees in 100% of the data sets, consistent with the findings of Khakurel et al. (in press). Interestingly, the MkVP+G model was divided between larger and smaller trees compared to the Mk model, with only 35% of the trees being larger. Figure 2B shows the tree length plotted for two data sets, of Hyaenodontidae (Egi et al., 2005) and Proboscideans (Shoshani et al., 2006), respectively. This is to highlight, that while there are some general trends, models still behave differently depending on the data set. It is worth noting that the Shoshani et al. (2006) data set (Figure 2B (i)) is the larger of the two, both in terms of number of taxa and characters. The influence of different models on tree length tended increase with larger data sets, both in terms of taxa and character number see supplementary fig. S1.

**Figure 2:**
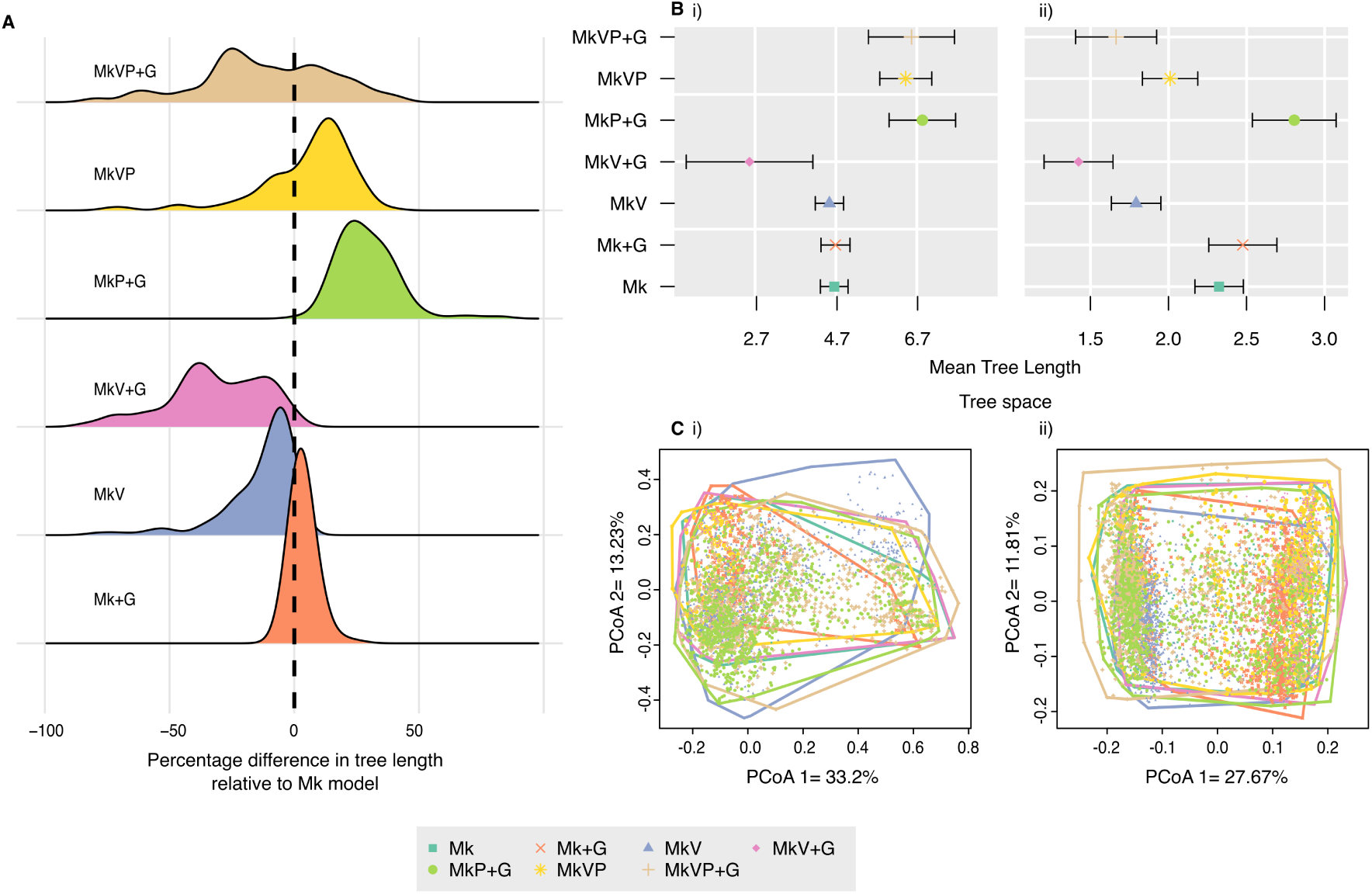
Analysis from 114 data sets under the 7 different models Mk, MkV, MkV+G, Mk+G, MkVP, MkVP+G, MkP+G. **A**, the changes in mean tree length of the posterior inferred using each model relative to the Mk model. **B**, the tree length calculated for each model for two different data sets from Egi et al. (2005) (Hyaenodontidae) and Shoshani et al. (2006) (Proboscideans), respectively. **C**, the tree space of the same two data sets as for B.

Figure 2C shows the tree space for the same two data sets. Using the permuted *p*-values estimated from the pairwise distances using Robinson-Foulds, we found that for both data sets the majority of models occupied a different tree space, i.e., differences in topology were significant. For the data set from Egi et al. (2005), trees inferred using MkV, MkV+G and Mk+G models grouped in a similar tree space, whereas all other models occupied different spaces. Whereas for the data set from Shoshani et al. (2006), we found two separate groupings, one of trees inferred using the Mk+G and MkV models, and the other an overlap between the MkV and MkV+G models. These results highlight that, not only do the substitution models have an impact on key parameter estimates, but this impact is not uniform across data sets.

### 4.2 Assessing the Performance of Model Adequacy and Model Selection Methods for Morphological Data

#### 4.2.1 Candidate Test Statistics for Morphological data

We explored the use of six test statistics for morphological models. The desired characteristic of test statics considered here, is their ability to indicate the adequacy of a particular model while also pointing out the inadequacy of another, i.e., we want the effect size of the correct model to be consistently around zero, while being far from zero for the incorrect models. We will focus on the results from both hyena and elephant data sets simulated under the MkV+G and MkVP+G models. We carried out the same investigation on data sets simulated under the MkV and MkVP models and reached the same conclusions, see Fig. S6–S8. The data test statistics, shown in Fig. 3, Grower’s coefficient and Generalized Euclidean Distance, both show a similar pattern. For the unpartitioned models there is no discernible preference for a given model. That is, they all fall within a similar range of effect sizes. For data simulated under a partitioned model, there was a stronger separation of effect sizes, where all the partitioned models are closer to zero and fall within a similar range. This pattern is more consistent for Gower’s Coefficient, suggesting it’s potential use as a test statistic. Neither of the inference based test statistics, shown in Fig. 4, show any strong or meaningful separation of effect sizes. Meaning, there is no preference for any of the models and it is unclear what explains this pattern. As for the mixed test statistics, consistency index and retention index, shown in Fig. 5, there is a similar pattern to that of the data based test statistics, however, with the differences in effect sizes between models being more pronounced. In order to quantify these results, we focused on three key features, (i) the variance in effect sizes for the correct model, meaning the total range of effect sizes for a given test statistic with the correct model, (ii) how incorrect models preformed, meaning the total range of effect sizes for a given test statistics across all models and, (iii) how easily we could differentiate between adequate and inadequate models by calculating the number of models which fall into the correct model effect size (ES) range. A numerical summary of these results can be found in Table 2 and 3. Consistency index and retention index demonstrated the best performance of these three aspects, with the correct models being consistently close to zero, incorrect models having larger ES values, and the fewest number of models on average falling within the correct model effect size range. While Grower’s coefficient also seems promising, the difference in effect sizes is less than that of the mixed test statistics. As such, in the empirical analyses we relied solely on the mixed test statistics, the consistency and retention indices.

**Figure 3:**
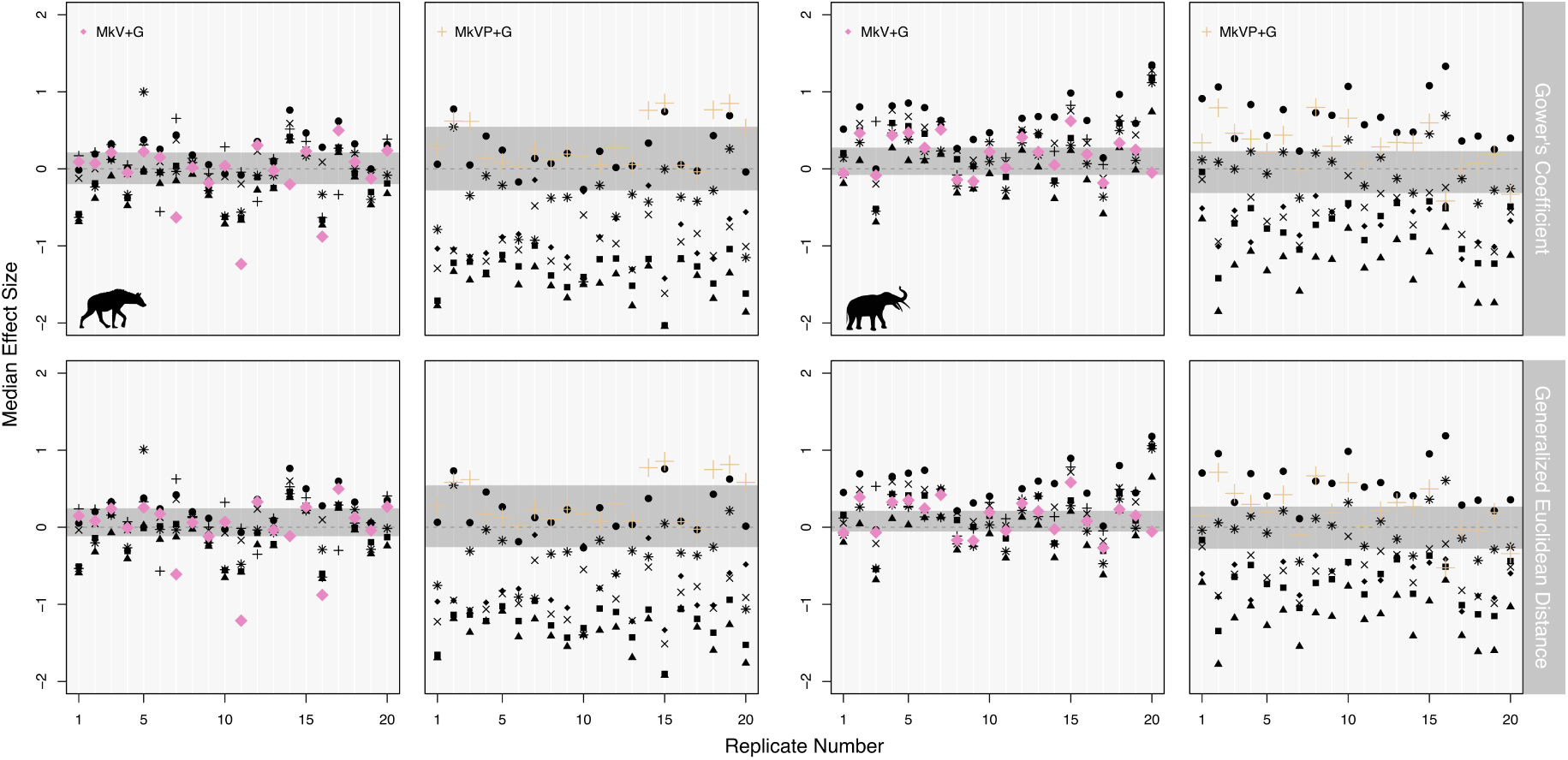
Validation of the data based test statistics. Plots show the output from each simulated data set with 20 replicates for each test statistic. The coloured points indicate the correct model, with the grey horizontal bar marking the range of effect sizes calculated for the correct model. ▪ = Mk, ✕ = Mk+G, ▴ = MkV, ◆ = MkV+G, ✳ = MkVP, ● = MkP+G, and + = MkVP+G

**Figure 4:**
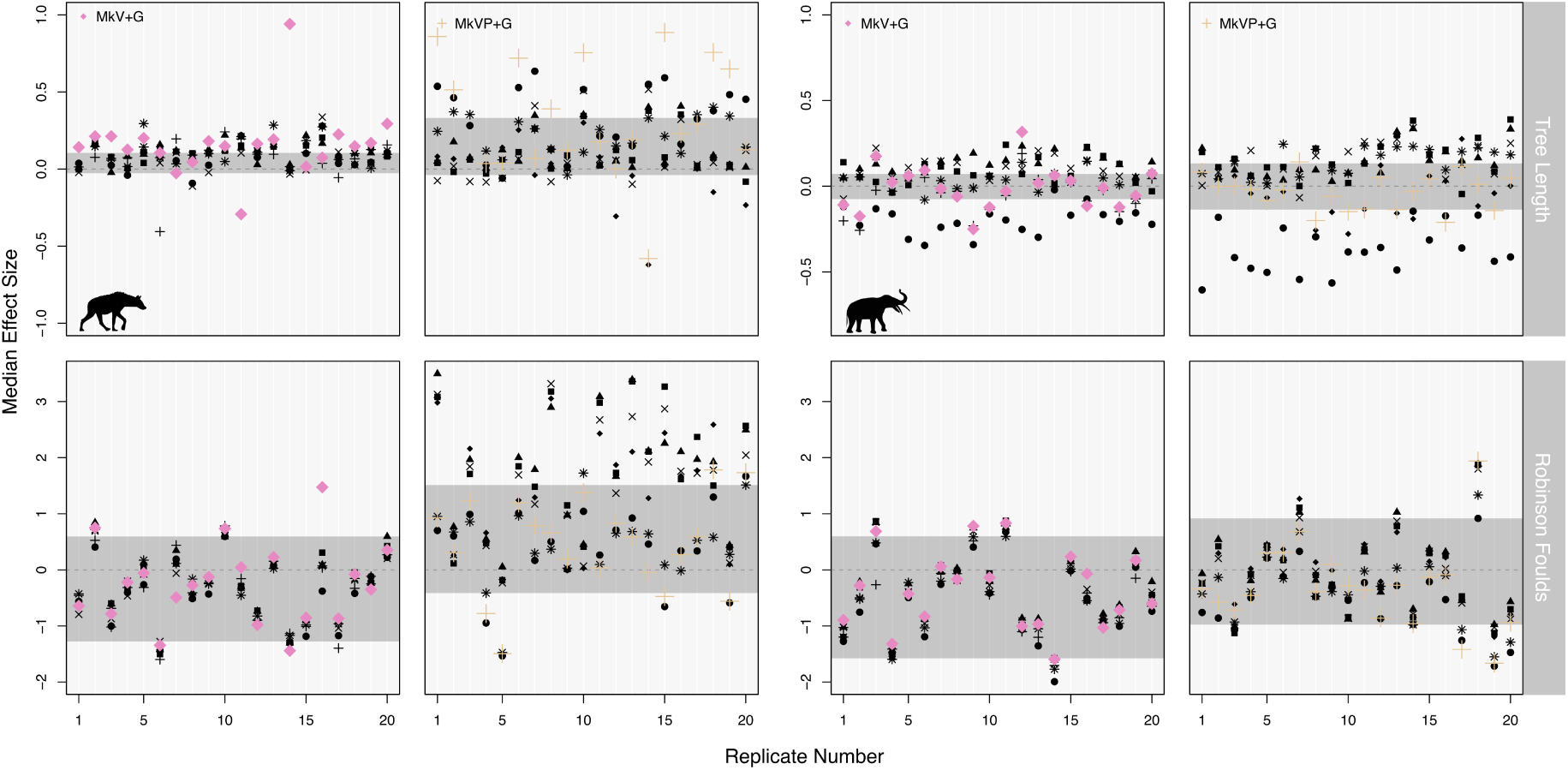
Validation of the inference based test statistics. Plots shows the output from each simulated data set with 20 replicates for each test statistic. The coloured points indicate the correct model with the grey horizontal bar marking the range of effect sizes values calculated for the correct model. ▪ = Mk, ✕ = Mk+G, ▴ = MkV, ◆ = MkV+G, ✳ = MkVP, ● = MkP+G, and + = MkVP+G

**Figure 5:**
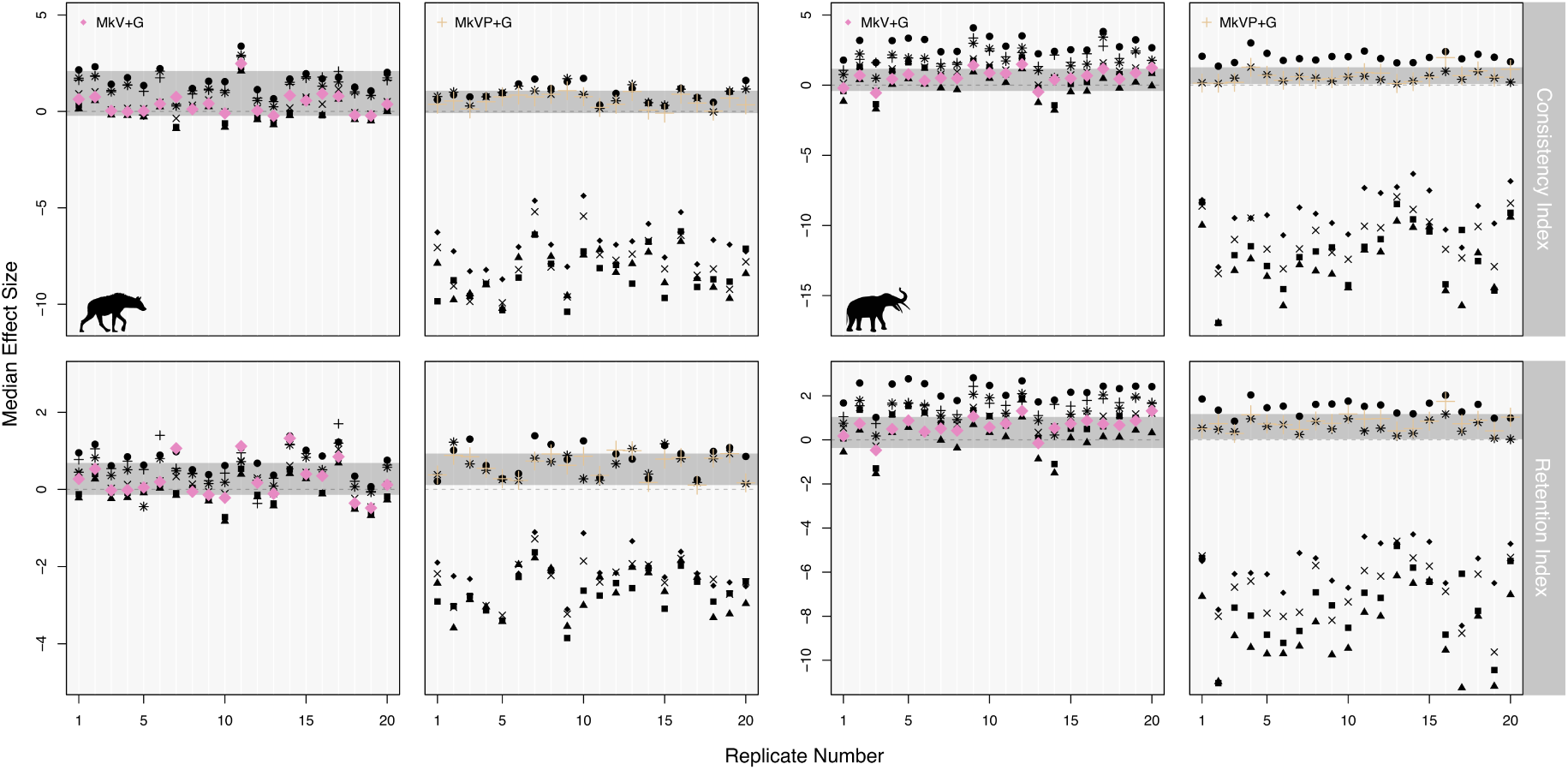
Validation of the mixed test statistics. Plots shows the output from each simulated data set with 20 replicates for each test statistic. The coloured points indicate the correct model with the grey horizontal bar marking the range of effect sizes calculated for the correct model. ▪ = Mk, ✕ = Mk+G, ▴ = MkV, ◆ = MkV+G, ✳ = MkVP, ● = MkP+G, and + = MkVP+G

**Table 2:** Validation of test statistics from the simulated hyena data sets. Correct ES gives the total range of effect sizes for a given test statistics with the correct model. Overall ES gives the total range of effect sizes for a given test statistic across all models. Num in Correct gives the number of models which fall into the Correct ES range. Num in Correct only looks at incorrect models, which means the maximum value here can be 6. GC = Gower’s coefficient, GED = generlized euclidean distance, TL = tree length, RF = Robinson Foulds, CI = consistency index, and RI = retention index. Consistency index and retention index have the largest overall ES range with, on average the fewest models falling in the same range as that of the correct model.

| Model | Test Statistic | Correct ES | Overall ES | Num in Correct |
| --- | --- | --- | --- | --- |
| MkV+G | GC | 1.7 | 2.2 | 5.7 |
|  | GED | 1.7 | 2.2 | 5.6 |
|  | TL | 1.2 | 1.4 | 6 |
|  | RF | 2.9 | 3.1 | 5.9 |
|  | CI | 2.7 | 4.3 | 5 |
|  | RI | 1.8 | 2.5 | 5.6 |
| MkVP+G | GC | 0.9 | 2.9 | 1 |
|  | GED | 0.9 | 2.8 | 1 |
|  | TL | 2.8 | 2.8 | 6 |
|  | RF | 3.3 | 5.0 | 4.1 |
|  | CI | 1.2 | 11.7 | 1.40 |
|  | RI | 1.1 | 5.3 | 1.6 |

**Table 3:** Validation of test statistics from the simulated elephant data sets. Correct ES gives the total range of effect sizes for a given test statistics with the correct model. Overall ES gives the total range of effect sizes for a given test statistic across all models. Num in Correct gives the number of models which fall into the Correct ES range. Num in Correct only looks at incorrect models, which means the maximum value here can be 6. GC = Gower’s coefficient, GED = generlized euclidean distance, TL = tree length, RF = Robinson Foulds, CI = consistency index, and RI = retention index. Consistency index and retention index have the largest overall ES range with, on average the fewest models falling in the same range as that of the correct model.

| Model | Test Statistic | Correct ES | Overall ES | Num in Correct |
| --- | --- | --- | --- | --- |
| MkV+G | GC | 0.8 | 2.0 | 4.1 |
|  | GED | 0.9 | 1.9 | 4.7 |
|  | TL | 0.7 | 0.7 | 5.7 |
|  | RF | 2.4 | 2.9 | 5.5 |
|  | CI | 2.0 | 5.9 | 2.8 |
|  | RI | 1.8 | 4.3 | 3 |
| MkVP+G | GC | 1.2 | 3.2 | 2.1 |
|  | GED | 1.2 | 4.0 | 2.95 |
|  | TL | 0.3 | 1.0 | 2.95 |
|  | RF | 3.7 | 3.7 | 5.95 |
|  | CI | 1.9 | 20.0 | 1.45 |
|  | RI | 1.5 | 13 | 1.6 |

#### 4.2.2 Model Adequacy vs. Model Selection

Here we compared the use of model adequacy and model selection using simulated data sets. To reiterate, unlike model selection, model adequacy approaches do not rank potential models in the same way, indicating that one model is the best. Therefore, for any given data set, if multiple models are investigated, as was the case here, several models may be adequate according to a particular test statistic. We will focus on the same 4 data sets as in section 4.2.1.

In the above section, to identify appropriate test statistics, we focused on the pattern of median ES values. When considering individual replicates we required more information than just the median ES value to determine the adequacy of a model for a given data set. Using this value alone makes it difficult to determine a model’s adequacy unless the median value is zero. We explored the use of upper and lower quartiles, and minimum and maximum limits and found the latter to be the more informative approach for identifying a model’s adequacy. We propose that if the minimum and maximum limits pass through zero, this would indicate that the model is adequate using our chosen test statistics. Following this criteria, we could quantify the percentage of simulation replicates where the model was deemed adequate/inadequate. Table 4 shows the percentage of times a model met the above described criteria using the consistency index and the retention index.

**Table 4:** The percentage of times a model was found to be adequate across all replicates using consistency index (CI) and retention index (RI) as tests statistics. In order for a model to be considered adequate the effect sizes need to meet the criteria put forward here, where the range of minimum and maximum values contain zero.

| Sim Model | Data set | Test Statistic | Mk | Mk+G | MkV | MkV+G | MkP+G | MVP | MkVP+G |
| --- | --- | --- | --- | --- | --- | --- | --- | --- | --- |
| MkV+G | Hyena | CI | 100% | 95% | 100% | 100% | 95% | 95% | 95% |
| MkV+G | Hyena | RI | 100% | 100% | 100% | 100% | 100% | 100% | 100% |
| MkVP+G | Hyena | CI | - | - | - | - | 100% | 100% | 100% |
| MkVP+G | Hyena | RI | 50% | 65% | 45% | 75% | 100% | 100% | 100% |
| MkV+G | Elephant | CI | 100% | 100% | 100% | 100% | 40% | 85% | 80% |
| MkV+G | Elephant | RI | 100% | 100% | 100% | 100% | 70% | 100% | 100% |
| MkVP+G | Elephant | CI | - | - | - | - | 100% | 100% | 100% |
| MkVP+G | Elephant | RI | - | - | - | - | 100% | 100% | 100% |

Model selection produced surprising results. We consistently found support for partitioned models, regardless of the model used to simulate the data. Table 5 shows the percentage of times a model was chosen as the best model according to Bayes factors. For this reason, using Bayes factors is not a reliable approach for deciding between partitions with morphological data, at least not using the standard approach we applied to partition characters, i.e., by the maximum observed state number (see the Discussion for a full explanation).

**Table 5:** Models chosen using Bayes factors and the marginal likelihoods. Cells show the percentage of times a model was selected across the 20 replicates from each simulation set up. The dashed line indicates the model was never selected.

| Model | Data set | Mk | Mk+G | MkV | MkV+G | MkP+G | MVP | MkVP+G |
| --- | --- | --- | --- | --- | --- | --- | --- | --- |
| MkV+G | Hyena | - | - | - | - | 5% | 15% | 80% |
| MkVP+G | Hyena | - | - | - | - | 5% | 30% | 65% |
| MkV+G | Elephant | - | - | - | - | - | - | 100% |
| MkVP+G | Elephant | - | - | - | - | - | - | 100% |

### 4.3 Analysis of Empirical data

We then applied PPS with the newly validated test statistics to 8 empirical data sets. This allowed us to answer our main question: are current morphological models adequate for empirical data? Of the 8 data sets, 5 had at least one model that was adequate. Fig. 6 shows the effect sizes from 4 data sets (see also supplementary Fig. S9). The MkVP+G model was found to be adequate for all 5 data sets. Of those 5 data sets, 4 also fit an MkVP model. We found the MkP+G model to be adequate for 3 data sets. For one of the data sets, Fig.6D, we found all models apart from the MkP+G model to be adequate. We do not see any clear pattern in terms of adequate models, with respect to the size of the data sets, i.e., number of taxa, characters, or state number. This suggests that these variables are not informative when choosing a model. For the two largest data sets, in terms of taxa, we did not find any models to be adequate. These data sets had 40 taxa (Shoshani et al., 2006) and 50 taxa (Tomiya, 2011). However, no models were adequate for a third data set with only 25 taxa (Schoch and Sues, 2013). Table. 6 shows the *p*-values calculated for consistency index and retention index for the same data sets as in Fig. 6. Values below 0.025 and above 0.975 are considered to be significant. This would indicate that the simulated data is significantly different from the empirical data, and that the model does not capture the underlying data generating processes and therefore is not adequate for that data set. Results using effect sizes and *p*-values agree on the same models for all data sets. There is one instance when there is a disagreement using retention index. For the data set from Egi et al. (2005), the Mk+G model was accepted using the threshold that we defined for effect sizes and rejected using *p*-values. Both metrics rejected the model according to consistency index, however, so the Mk+G was ultimately rejected using both approaches.

**Figure 6:**
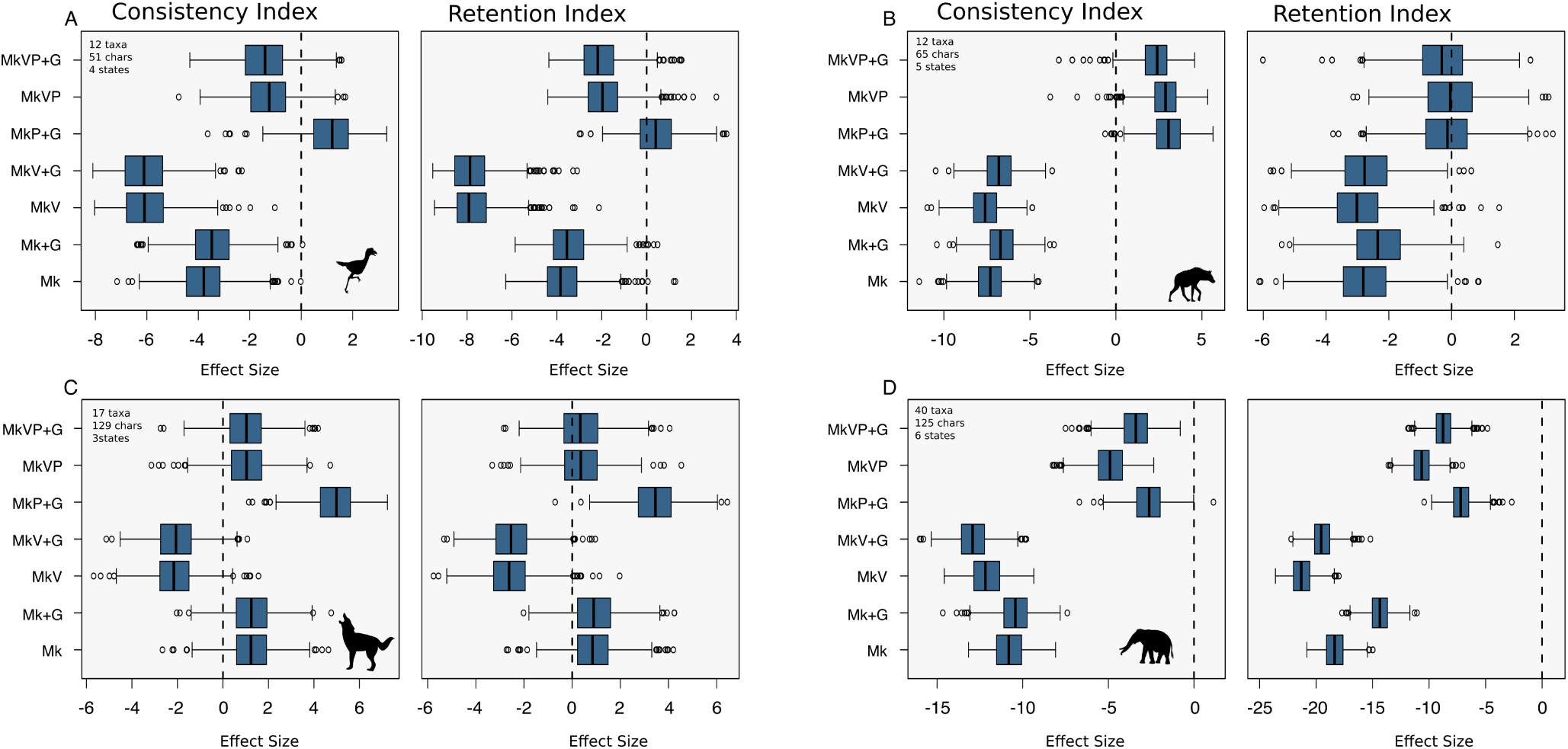
Effect sizes for four empirical data sets for the consistency index and retention index. The dashed black line is at zero is there to help identify adequate models. The data sets are taken from (Agnolin, 2007), (Egi et al., 2005), (Bourdon et al., 2009) and (Shoshani et al., 2006), respectively.

**Table 6:** Posterior *p*-values from the empirical analyses. CI refers to consistency index and RI to retention index. Values below 0.025 and above 0.975 are considered to be significant. This would indicate that the simulated data is significantly different than the empirical data and that the model is not adequate for that data set. The results here agree with those produced using effect sizes. See Table 4.

|  | Agnolin |  | Egi |  | Bourdon |  | Shoshani |  |
| --- | --- | --- | --- | --- | --- | --- | --- | --- |
| Model | CI | RI | CI | RI | CI | RI | CI | RI |
| Mk | 0 | 0.003 | 0 | 0.005 | 0.8895 | 0.812 | 0 | 0 |
| Mk+G | 0.001 | 0.004 | 0 | 0.006 | 0.898 | 0.8235 | 0 | 0 |
| MkV | 0 | 0 | 0 | 0.005 | 0.019 | 0.011 | 0 | 0 |
| MkV+G | 0 | 0 | 0 | 0.006 | 0.033 | 0.01 | 0 | 0 |
| MkP+G | 0.835 | 0.659 | 0.994 | 0.446 | 1 | 0.999 | 0.001 | 0 |
| MkVP | 0.105 | 0.041 | 0.992 | 0.483 | 0.859 | 0.655 | 0 | 0 |
| MkVP+G | 0.095 | 0.034 | 0.974 | 0.376 | 0.848 | 0.6245 | 0 | 0 |

## 5 Discussion

Understanding morphological evolution is an extremely difficult task. Within palaeobiology we rely on a small number of relatively simple models to describe this complex process (Wright, 2019). Until now, the impact of these different substitution models on parameter estimates was not well understood. Our analysis on the influence of these models using empirical data sets, focusing on tree length and topology, demonstrates that different models can produce contrasting reconstructions of the evolutionary history of a group, emphasising the importance of model choice (Fig. 2). Although the impact of models on parameter estimates is not uniform across data sets, the most consistent pattern we observe is whether or not the data is partitioned.

### 5.1 Partitioned models

In all the partitioned models explored here, traits were partitioned based on the number of character states. This is a practical approach, both in terms of the biology and the way in which the characters tend to be coded. We found that for all but two data sets, the unpartitioned models produced smaller trees. To further investigate the cause of this, we ran an analysis using a binary data set and increased the Q-matrix size from 2-5. The objective here was to mirror what happens when we have characters with a lower number of observed states than the maximum number of states in the matrix. For example, placing binary characters in a partition with a maximum of 5 character states. We show that as the size of the Q-matrix increases tree length gets smaller (Fig. S10). The effect of partitioning that we observe on empirical estimates of tree length, is therefore a direct result of how morphological data is typically partitioned (see also Equations 8 and 9 below). Characters are partitioned by maximum number of observed states, e.g., binary characters are all together in one partition and assigned to a rate matrix of size 2, characters with 3 states are assigned to a rate matrix of size 3 and so on. For unpartitioned models, however, all of the characters will be in a single Q-matrix that is the size of the maximum number of observed states across the whole data set. This means that for a given branch length *v*, under a model that assumes there are *n* states, for characters where we observe *<n* states (e.g., a binary character in a rate matrix of size 5), the probability of observing no change will be underestimated. Similarly, the probability of observing a given change will also be lower if there are more (unobserved) possible states. Both cases will result in shorter branch lengths. Partitioning morphological data by character state number is a practical approach, however, this requires making an assumption that we know the number of states for each character, when in reality we might not. For molecular data of course, this is not something we need to consider, as we know there are four nucleotides. By assuming we know the number of states, based on the number of observed states, we may be biasing our results. The effects of whether or not a data set is partitioned are considerable in terms of parameter estimates. As such, it is important to consider how the data is being partitioned and whether or not it makes biological sense for your data set to do so.

Here we focused exclusively on partitioning by character state. This is the most common partitioning scheme and is even a default in some phylogenetic software programs, for example BEAST2 (Bouckaert et al., 2019), and MrBayes (Ronquist et al., 2012). Yet this is not the only way that data could be partitioned. A researcher could partition the data based on different anatomical regions, or based on subsets of anatomical, ecological or behavioural traits. Thus one may need to decide between various partitioning schemes or no partitioning at all. To date, model selection is regarded as the gold standard for choosing between substitution models and partition schemes (Xie et al., 2011). Within a Bayesian framework, comparing marginal likelihoods has been shown to be effective for choosing between partition schemes with molecular data. Our results, however, show that for morphological data, model selection consistently selects a partitioned model, regardless of the model used to simulate the data. This result can be explained by taking into account how partitioning morphological data effects the likelihood calculation, importantly how it effects the transition probabilities and the stationary frequencies.

For example, assume you have a tree consisting of two tips, one with discrete state 0 and the other with discrete states 1, as shown here.

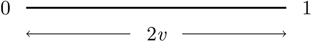

The tips share a common ancestor *v* time units in the past. The transition probability for this scenario under the Mk model is calculated as:

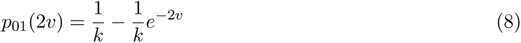

where *k* is the number of states. Further, the likelihood of this data is:

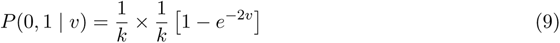

Here *k* would be set to 2 as we observe two states. However, in cases where there are other traits, some of which have a higher maximum observed state, *k* would increase., e.g., as happens in unpartitioned inference. Higher values of *k* would result in a lower likelihood. This change in likelihood is a direct result of the partitioning scheme. When partitioning molecular data, we do not change the size of the Q-matrix (*k*), which is why we do not see the same effects on the likelihood. Figure 7 shows the impact on the log likelihood of changing the size of the Q-matrix (*k*) along different branch lengths (*v*) for these two tips.

**Figure 7:**
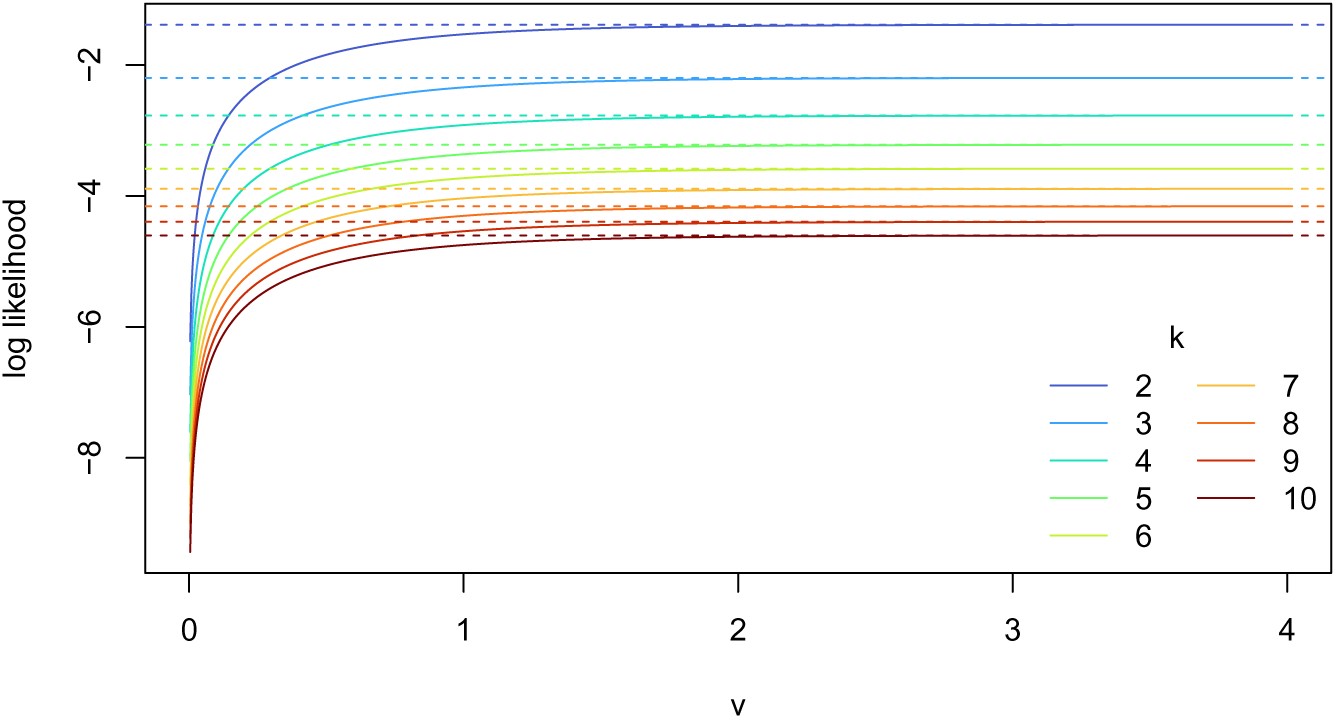
Log likelihoods calculated for different sizes Q-matrices (k) along as a function of branch lengths (*v*). The log likelihoods converge as *v* increases and the transition probability approaches the stationary frequencies.

To empirically demonstrate the impact of of partitioning by state space on the likelihood we ran two experiments. First, using an empirical binary morphological matrix we calculated the marginal likelihood under an unpartitioned MkV+G model increasing the Q-matrix size from 2-5. Supplementary figure. S12 shows the decrease in marginal likelihood as we increase the number of transition possibilities (Q-matrix size). We then wanted to investigate the impact of adding the “correct” partitions. Here, we used an empirical morphological matrix with a maximum of 6 states. We first calculated the marginal likelihood under an unpartitioned MkV+G model. We then created two partitions, one partition for all binary states and the second for all other states. Then we increased the number of partitions to three, with one for binary states, one for ternary states and kept all others in the third partition. This method of adding partitions was continued until there were 5 in total and all states were in the appropriately sized Q-matrix. Fig. S11 shows that the marginal likelihood increases as partitions are added to the model. This is expected, given Equations 8 and 9. This suggests that the results from model selection will not be indicative of any meaningful biological signal. For this reason, using model selection to differentiate between partitions for morphological data is not appropriate when the Q-matrix size varies.

### 5.2 Test Statistics

Overall, our results show that model adequacy, in particular PPS, currently offers the most effective way of identifying the most suitable model for morphological data. In addition, we demonstrate that PPS can reliably determine whether a given model is adequate or not. Understanding the absolute fit of available models can lend support to the use of model based phylogenetics for the analysis of morphological data. Here we carried out the first thorough investigation into the use of PPS with discrete morphological substitution models.

One of the most important aspects of PPS to consider is the choice of test statistics. As this was the first systematic application of PPS to discrete character data, we first validated available test statistics using simulations. We explored the use of 6 test statistics and ultimately found consistency index and retention index to be the most informative. Neither of the inference based test statistics we explored, Robinson-Foulds or tree length, were able to give a clear indication of model adequacy. In this context, Robinson-Foulds distance is used to quantify variance across the posterior distribution of trees, therefore reflecting topological uncertainty. Given that morphological data sets tend to be small, the uncertainty in topology may be high, regardless of the model used for inference (Barido-Sottani et al., 2020). The uninformativeness of tree length is more puzzling, since competing models have a clear impact on the estimated tree length. Tree length has also previously been shown to be a poor test statistic for molecular data (Duchêne et al., 2018). Both Gower’s coefficient and generalized euclidean distance did show some potential value as test statistics (Fig. 3), although the mixed test statistics, the consistency index and retention indices, were substantially better (Fig. 5, Tables 2–3). Having a test statistics specifically focused on the data would be favourable. Future studies could focus on alternative ways of including disparity metrics as test statistics. For example, we used the mean pairwise distance, perhaps looking at the sum of the variance or sum of the ranges could be more informative for model adequacy (Smith et al., 2023).

### 5.3 Practical Considerations

Importantly, our simulation study also allowed us to identify ways of reducing the overall computational costs. As with many Bayesian analyses, there can be a high computational costs associated with running a PPS analysis. To mitigate any unnecessary computation, we assessed the maximum number of simulation replicates required to reach stability in the mean effect sizes. By doing so, we were able to ensure that we were not running unnecessary replicates. Further, the most expensive part of running a PPS analysis comes from the inference of the simulation replicates. Based on our simulation study, we did not find any benefit to including inference based test statistics (tree length and Robinson Foulds, Fig.4), meaning this expensive step can be skipped. Taking both of these findings into account, the time and memory required to run a PPS analysis becomes a lot smaller. For example, when compared to a stepping stone analysis, we found PPS to take half the time per model.

From our simulation study, relying exclusively on the mixed test statistics, consistency index and retention index, we found that for all replicates, more than one model was adequate (Table 4). When interpreting these results it is important to remember simulated data is often “neater” than empirical data. In our simulation set up, all characters in a given matrix were simulated under the same model and the model extensions we used are not proposing conflicting statements about the underlying process. As such, it is not surprising that we found multiple models to be adequate for our simulated data. The choice of substitution model may have less impact on our simulated data, as the topology is easier to infer. For example, taking all simulation replicates of the simulated hyena data under an MkV+G model, the mean variance in tree length across the 7 different models was 0.74. In contrast, for the empirical data used as the basis for the simulations, the variance in tree length across models was 4.29 (Fig. 2B(i)). Our simulation study was valuable in determining which test statistics were sensitive to model choice under exemplar conditions, but it is not alarming that differentiating between similar models, i.e., all partitioned models, was not possible. Future work could investigate model adequacy when data is simulated under more complex models, e.g., generating matrices that contain conflicting characters associated with different models or topologies (Sansom et al., 2017; Weisbecker et al., 2023).

The results from our empirical data sets show a larger difference in the effect sizes for different models (Fig. 6). Based on our criteria of using the minimum and maximum effect sizes (after removing outliers) we determined that for 5 of data sets, at least one of the models tested here was adequate. This leaves the other 3 without a model being adequate. While initially this result may seem negative, in that no models were adequate, it is actually more reasonable than not. The expectation that all data sets would have a model available that fit would have been unrealistic, given the complexity of the data versus the simplicity of the models. Having a method which allows the researcher to detect the limits of available models is much more useful than picking the best out of a group of models without considering whether any of them fit. This result highlights the benefit of using such an approach. In the situation where no models are considered adequate for a data set, it would be up to the researcher to determine how to proceed. For instance, if the effect sizes are not markedly far from zero one may still opt to use a model, however, appreciating its limitations would be important before drawing any conclusions based on the inference results. It is also encouraging to see that the most complex model, the MkVP+G model, was identified as adequate for all 5 of the data sets for which we found an adequate model, indicating that we are moving in the right direction, in terms of our assumptions about the data generating processes. This strongly supports the above discussed rationale of partitioning the data based on character state, lending confidence to our biological interpretation of the evolution of the data.

Here we have demonstrated how PPS outperforms a model selection approach in several respects. Making this a standard approach in palaebiology would be beneficial to the field in allowing for a better appreciation of how well our models are performing. In this study we explored the use of 7 extensions of the Mk model, as they are the most commonly applied. This is not an exhaustive list of available models and there are a number of alternatives that further relax assumptions of the Mk model. For example, Wright et al. (2016) showed how relaxing the assumption of symmetrical probability of change between characters can improve model fit and phylogenetic estimation (Wright et al., 2016). Models including hidden states have also been proposed for morphological data (Tarasov, 2019). Such models can also be assessed using the workflow presented here, the only requirements being that the model can be used for both simulation and inference. There are also a number of models of continuous character evolution that are often used in phylogenetic comparative methods (Álvarez-Carretero et al., 2022; Hansen et al., 2022). While we did not explore these models here, there has been work previously carried out demonstrating their use with model adequacy (Slater and Pennell, 2014). We focused exclusively on discrete data as it remains the most widely used for tree inference. Our results also have implications for studies focused on divergence time estimation and ancestral state reconstruction. The same model validation can be applied before either of these types of analyses are carried out. Fossils are our only direct source of information about extinct taxa. Collection and character coding of fossils for phylogenetic analysis requires huge effort both in terms time and knowledge required. Ensuring that we are using the best available models can help provide confidence in our results and ask more complex questions with the data.

## 6 Conclusions

As model-based phylogenetic analysis gains prominence in paleobiology our study aimed to emphasise the importance of model choice, by demonstrating how different substitution models can impact inference results. We show that substitution model choice impacts estimates of both lengths and topology. By providing a workflow for PPS to validate models adequacy, researchers can gain insights into absolute rather than relative model fit, and can have more confidence in their choice of substitution model going forward. We show that, despite the arguably simplistic assumptions of available morphological models, they are often able to approximate the underlying generating processes of discrete morphological data sets. However, we also show that no single model fits all data sets examined here, so we recommend researchers use model adequacy to assess model fit as a first step in phylogenetic inference. Given the substantial taxonomic effort invested into collecting these data sets, the importance of utilizing accurate models cannot be overstated. Our work reinforces the significance of these considerations, particularly as fossil data remains the primary avenue for gaining a comprehensive understanding of evolutionary history in deep time.

## 7 Supplementary Material

All data sets used here were taken from previous studies and are available on GitHub (https://github.com/laumul/PPS_Morphology). The associated RevBayes tutorial is available here (https://revbayes.github.io/tutorials/pps_morpho/pps_data_morpho.html).

## 8 Acknowledgements

AMW was supported on NSF DEB-2045842. SH was supported by the Deutsche Forschungsgemeinschaft (DFG) Emmy Noether-Program (Award HO 6201/1-1 to S.H.) and by the European Union (ERC, MacDrive, GA 101043187). Views and opinions expressed are however those of the authors only and do not necessarily reflect those of the European Union or the European Research Council Executive Agency. Neither the European Union nor the granting authority can be held responsible for them. MRM was supported on NSF DEB-1754705. The majority of this research was conducted with high-performance computational resources provided by Friedrich Alexander University Erlangen-Nuremberg (https://hpc.fau.de).

## 9 Supplementary Information

**Figure S1:**
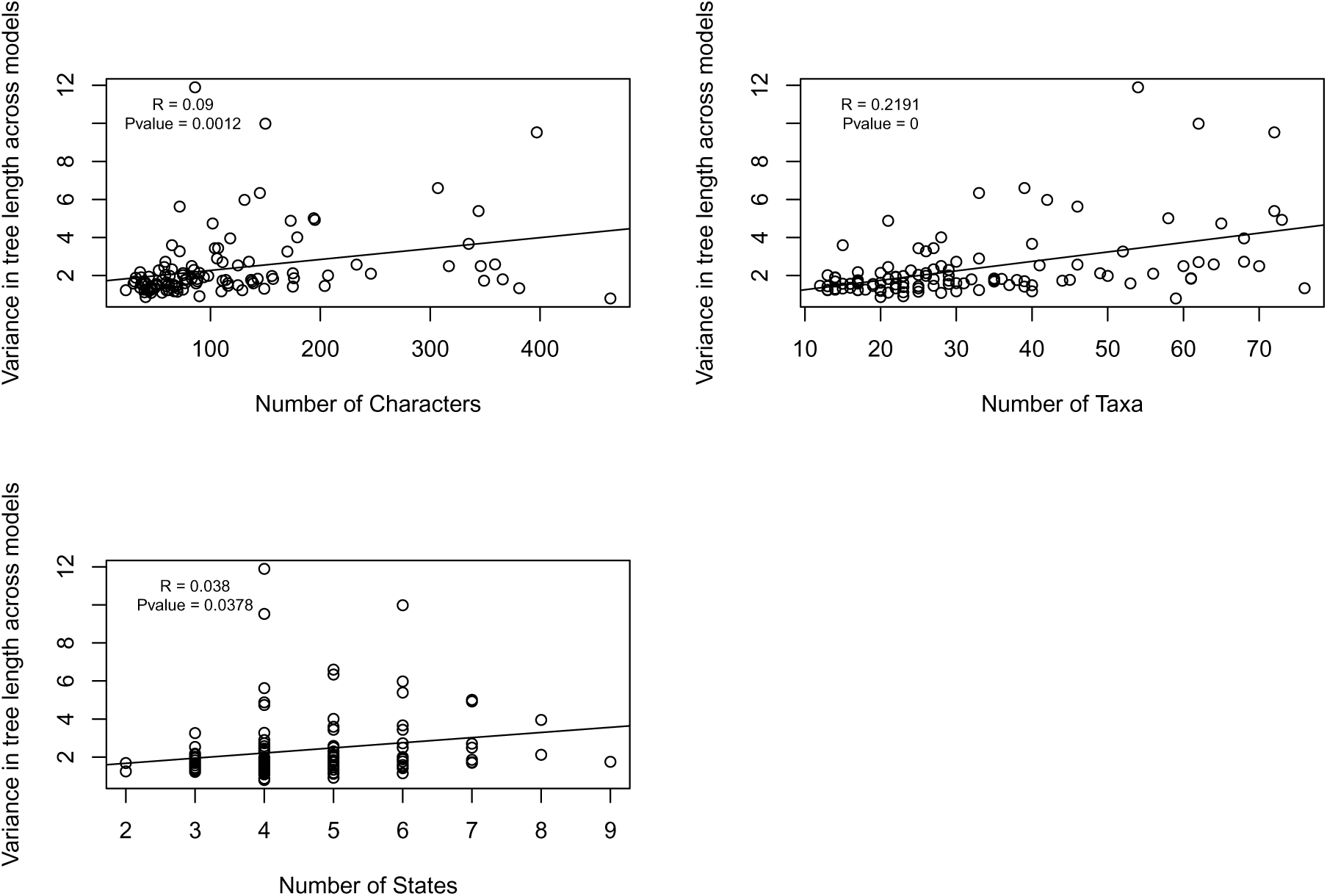
The relationship between the impact of different models on branch lengths and properties of the data sets. Variance between models was calculated by subtracting the smallest tree length from the largest tree length for each data set, irrespective of the model used for inference.

**Figure S2:**
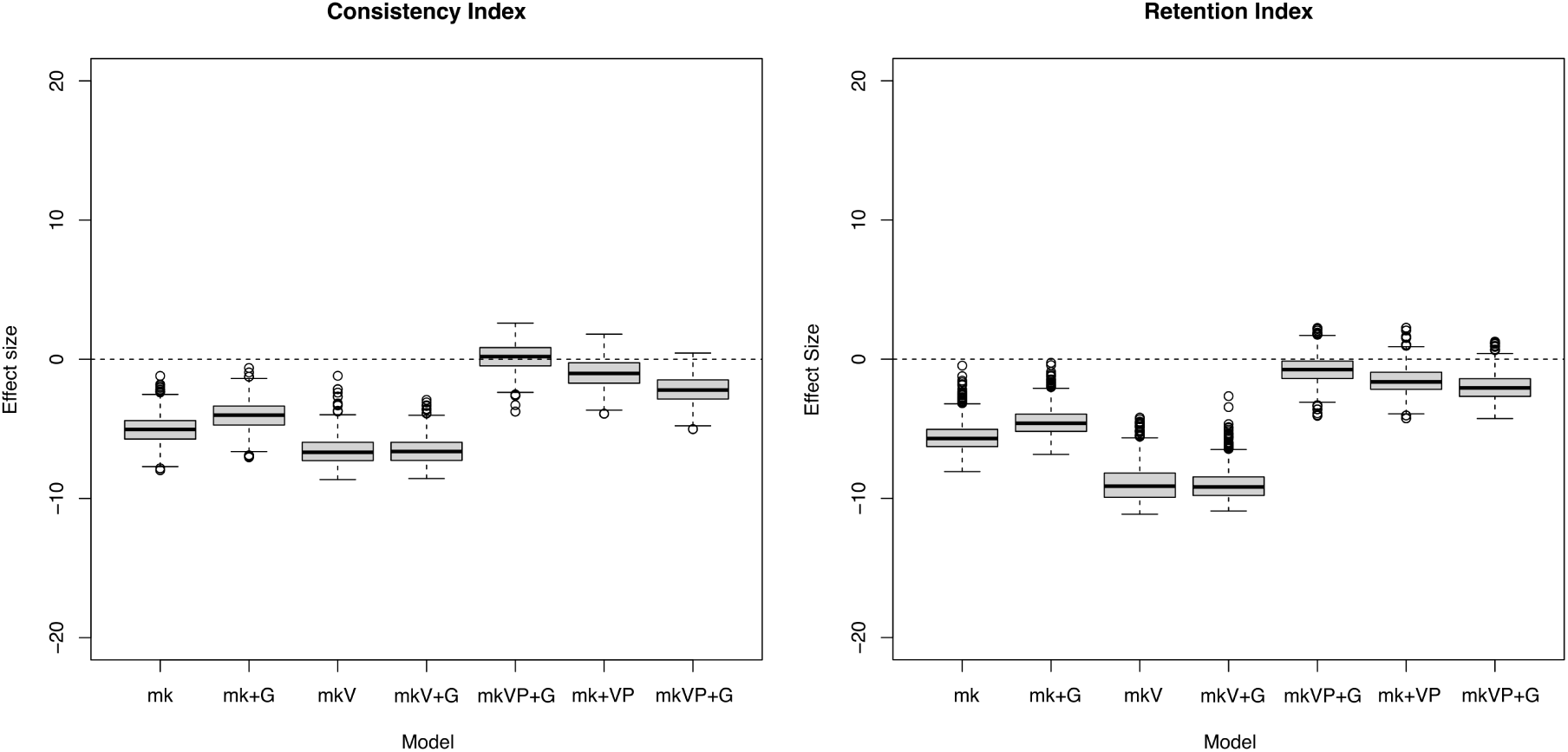
Effect Sizes calculated for the Agnolin data set using the entire posterior distribution. While the values are slightly different compared to those calculated using an MCMC tree, see fig. 6A, it determines the same models being adequate. This calculation also increased computation time significantly.

**Figure S3:**
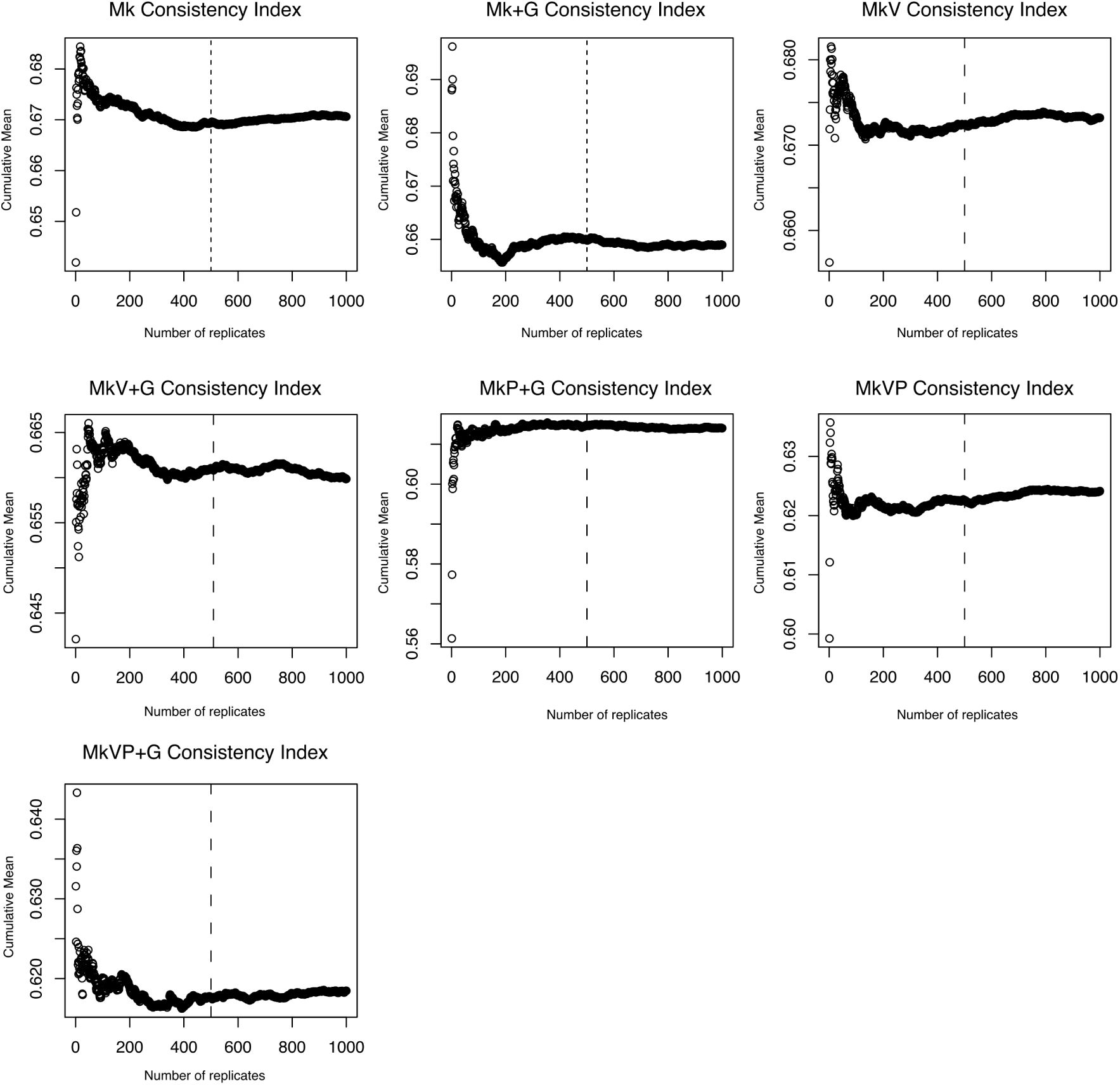
The cumulative means calculated for consistency index for one replicate of the simulated hyena data sets under the MkV+G model. This serves as a representative of all other replicates and test statistics which also showed the same pattern. The dashed line is at 500. After this point the line plateaus, representing that the variation of mean effect size is constant after that point.

**Figure S4:**
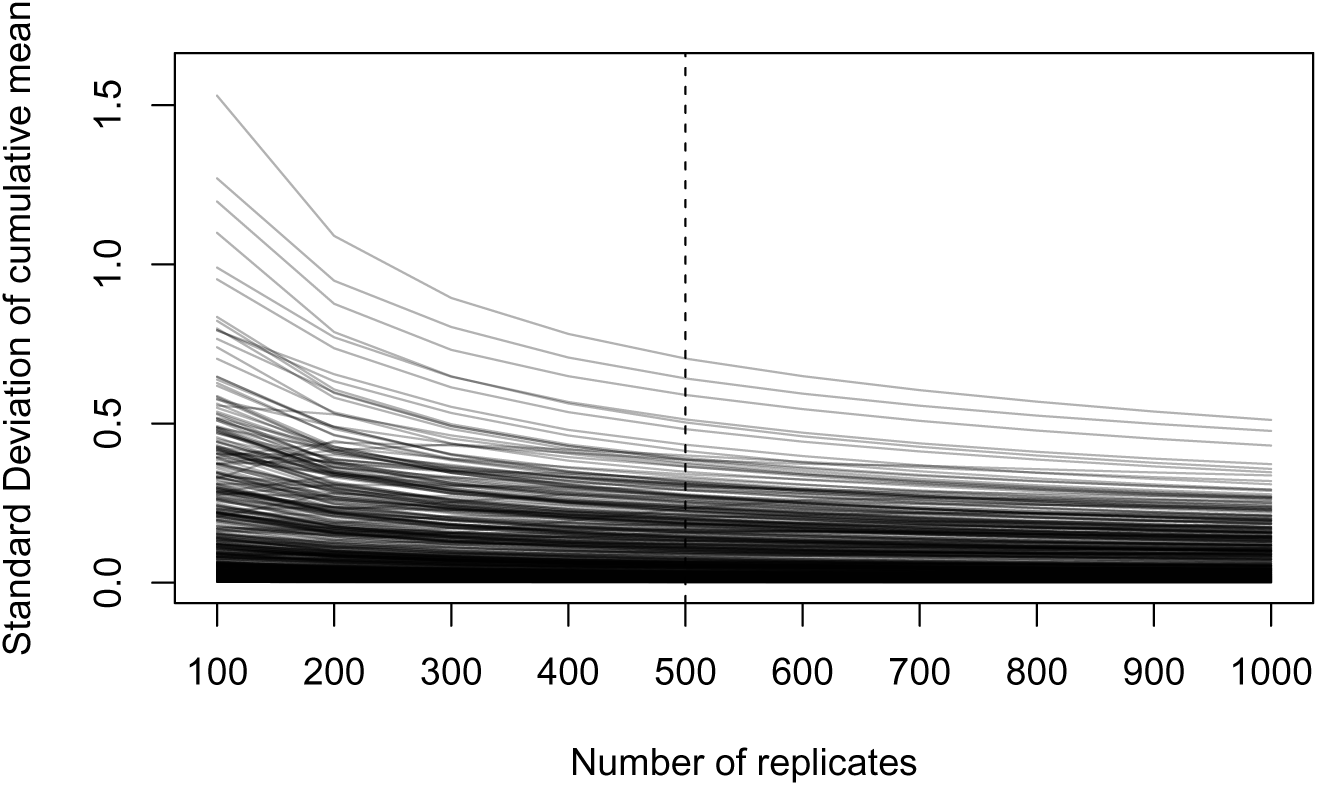
The standard deviation around the cumulative mean for all replicates of the simulated hyena MkV+G and test statistics (Gower’s coefficient, generalized euclidean distance, tree length, Robinson Foulds, consistency index, and retention index). The dashed line is at 500. After this point the lines plateaus, indicating that the mean will not change if more replicates are added.

**Figure S5:**
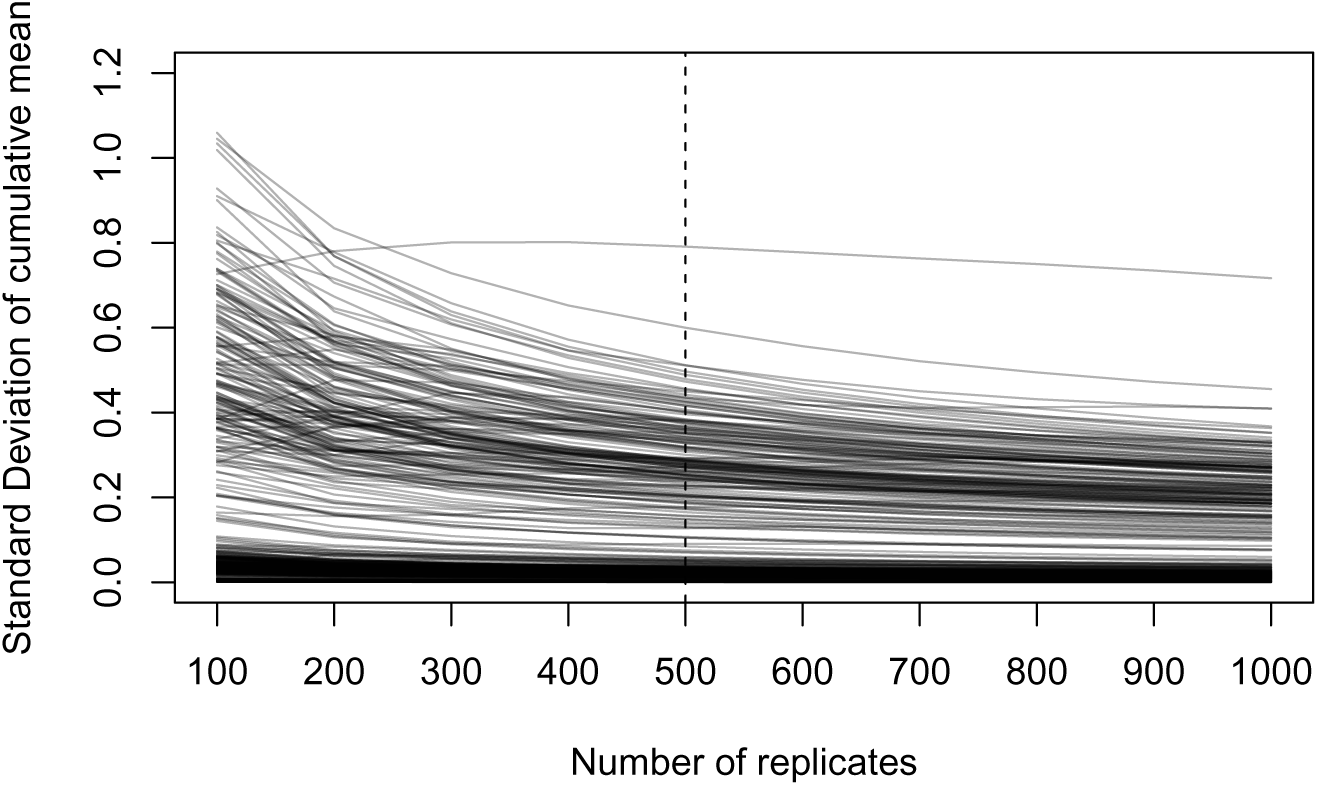
The standard deviation around the cumulative mean for all replicates of the simulated elephant MkV+G and test statistics (Gower’s coefficient, generalized euclidean distance, tree length, Robinson Foulds, consistency index, and retention index). The dashed line is at 500. After this point the lines plateaus, indicating that the mean will not change if more replicates are added.

**Figure S6:**
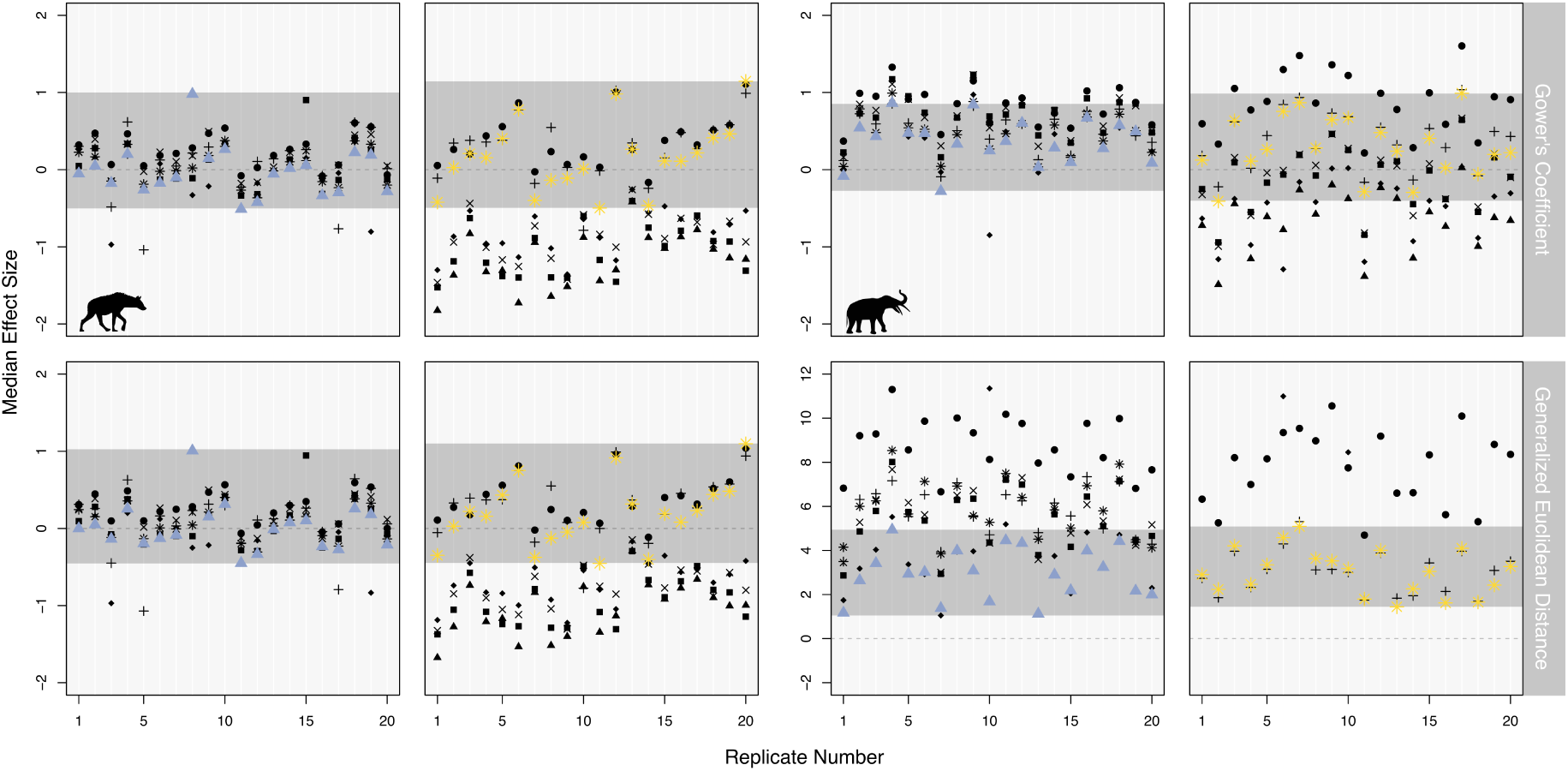
Validation of the data based test statistics. Plots show the output from each simulated data set with 20 replicates for each test statistic. The coloured points indicate the correct model, with the grey horizontal bar marking the range of effect sizes calculated for the correct model. ▪ = Mk, ✕ = Mk+G, ▴ = MkV, ◆ = MkV+G, ✳ = MkVP, ● = MkP+G, and + = MkVP+G

**Figure S7:**
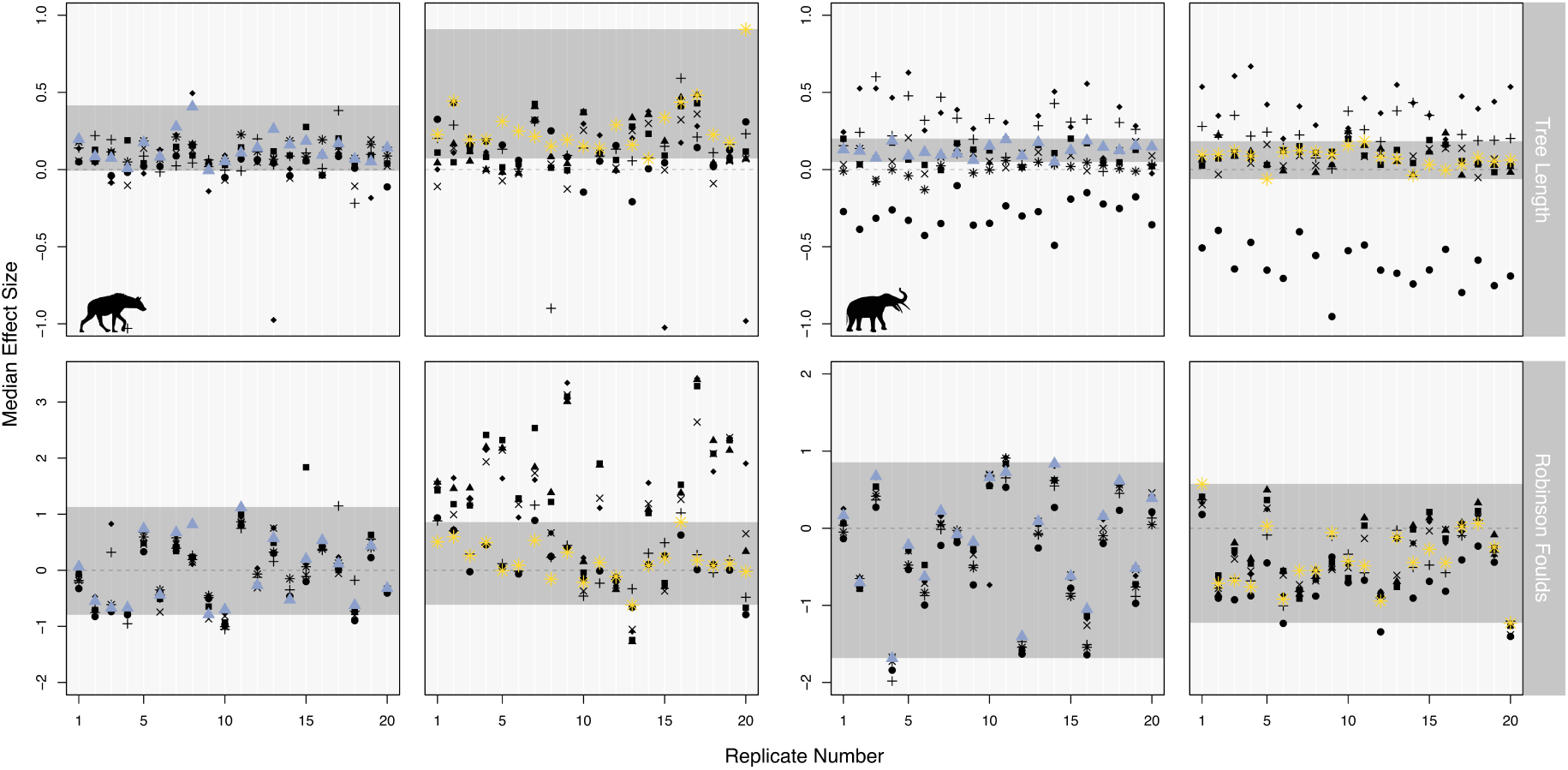
Validation of the inference based test statistics. Plots shows the output from each simulated data set with 20 replicates for each test statistic. The coloured points indicate the correct model with the grey horizontal bar marking the range of effect sizes values calculated for the correct model. ▪ = Mk, ✕ = Mk+G, ▴ = MkV, ◆ = MkV+G, ✳ = MkVP, ● = MkP+G, and + = MkVP+G

**Figure S8:**
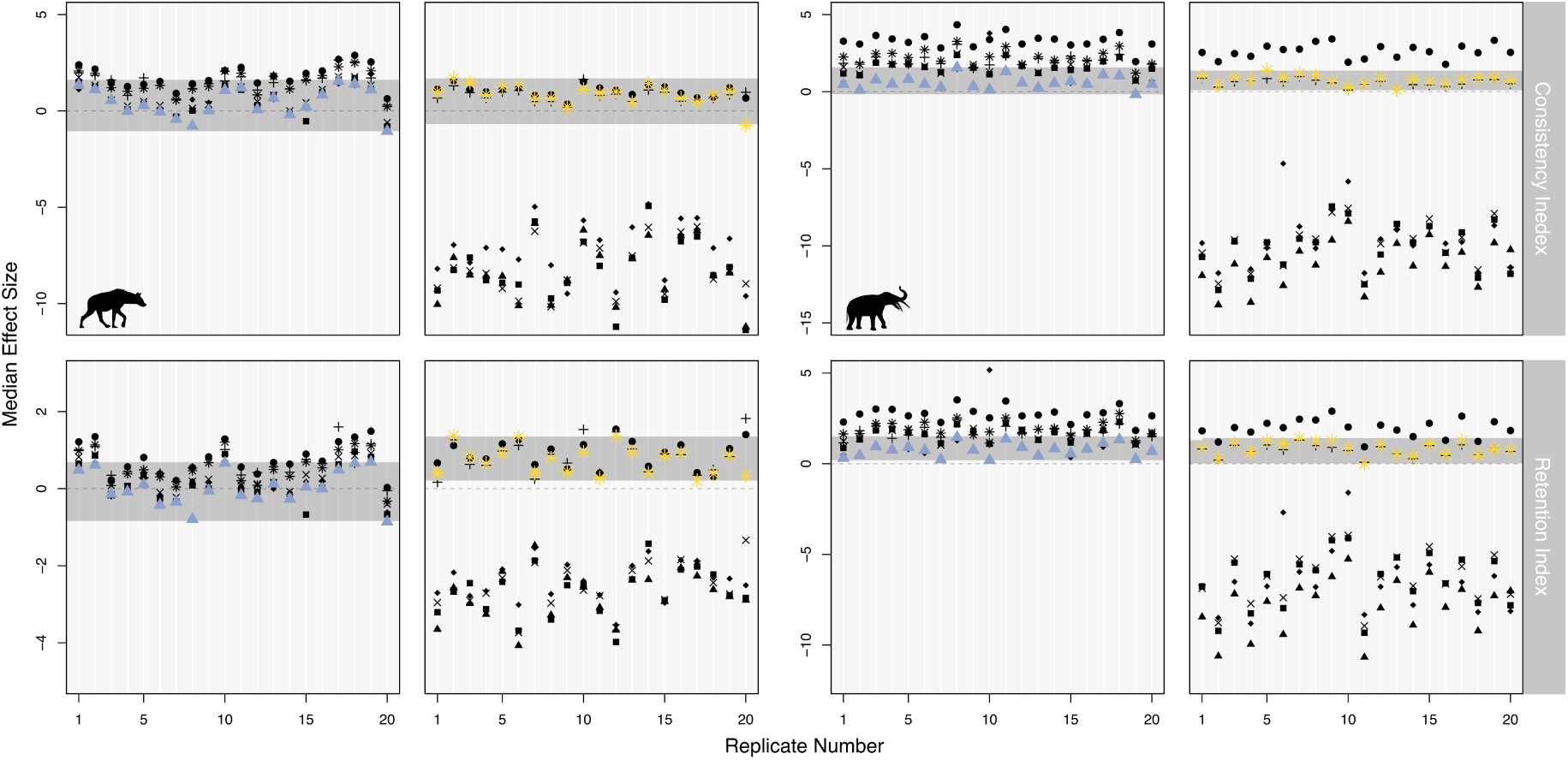
Validation of the mixed test statistics. Plots shows the output from each simulated data set with 20 replicates for each test statistic. The coloured points indicate the correct model with the grey horizontal bar marking the range of effect sizes calculated for the correct model. ▪ = Mk, ✕ = Mk+G, ▴ = MkV, ◆ = MkV+G, ✳ = MkVP, ● = MkP+G, and + = MkVP+G

**Figure S9:**
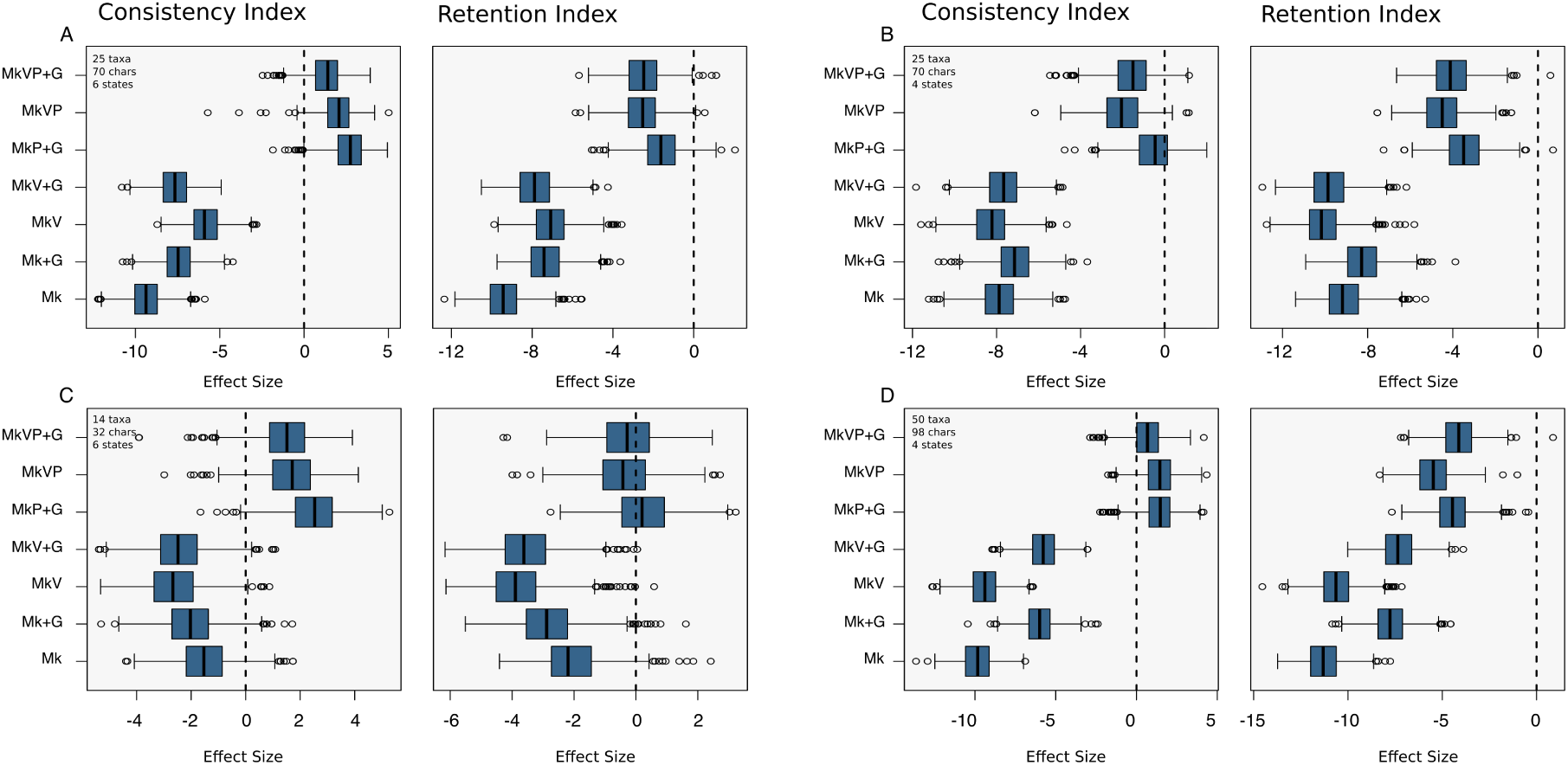
Results from four of the empirical data sets for consistency and retention index. The dashed black line is at zero is there to help identify adequate models. The data sets are taken from Archibald et al. (2001), Schoch and Sues (2013), Bloch et al. (2001), and Tomiya (2011) respectively

**Figure S10:**
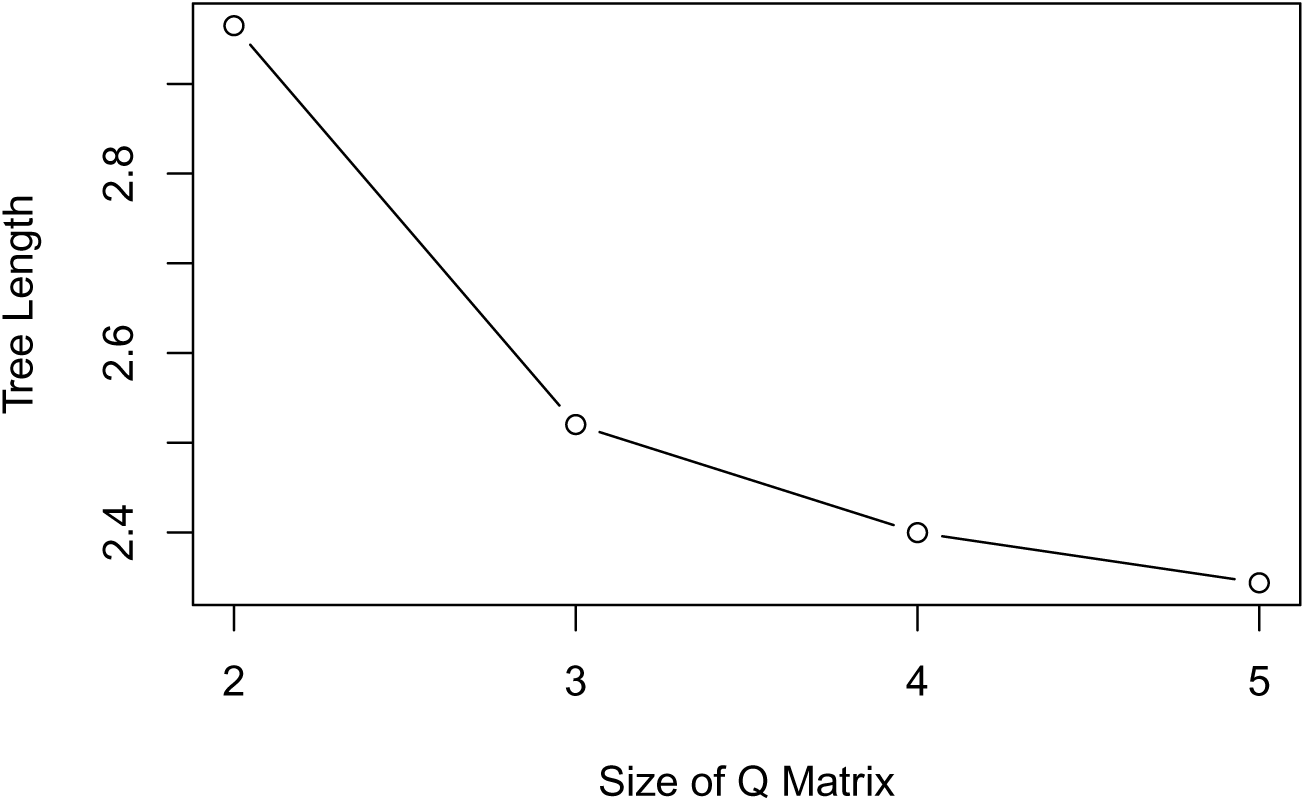
The impact of the Q-matrix size on tree length. Using a binary alignment the Q-matrix was increased from 2-5. The tree length becomes smaller as the Q-matrix increases in size.

**Figure S11:**
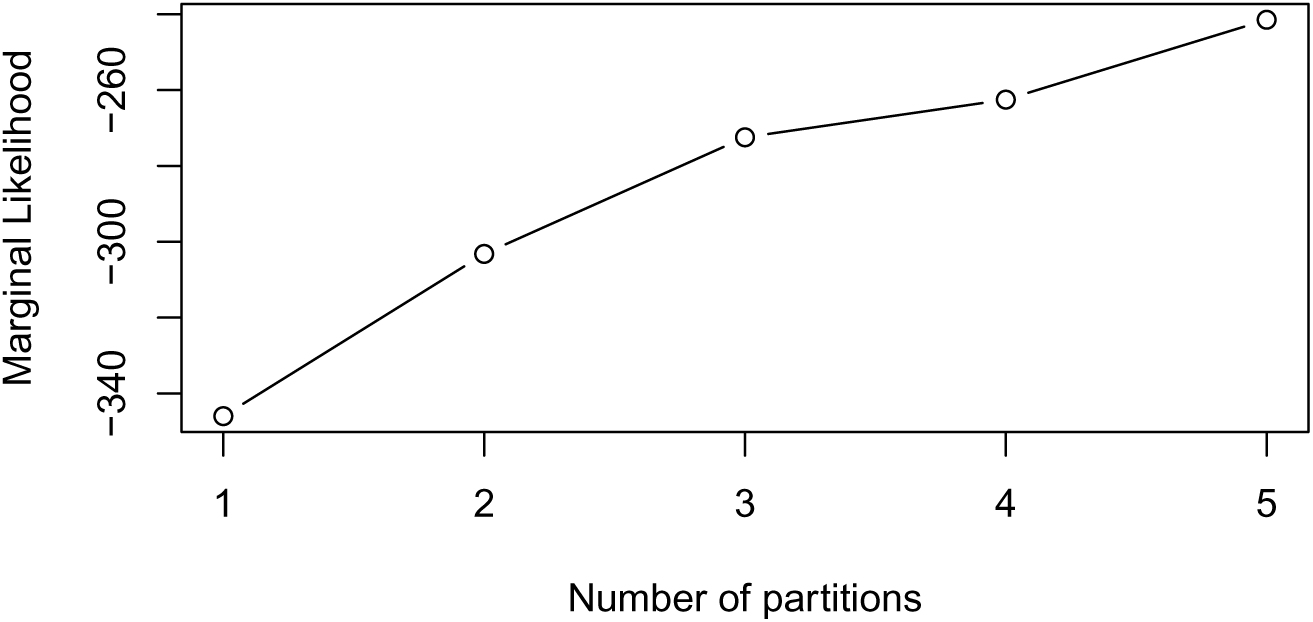
The effects of increasing partitions on the likelihood calculation. Using a data set with a maximum state size 6 the number of partitions was increased from 1 to 5. Where 1 was completely unpartitioned, 2 has one partitions for binary characters with all others in the other partition, 3 has one partition for binary, one partition for tertiary and all others in the third partition, and so on until all characters are in the correct partition with 5 partitions. As the number of partitions increase, and characters are added to a Q-matrix of the correct size, the likelihood increases.

**Figure S12:**
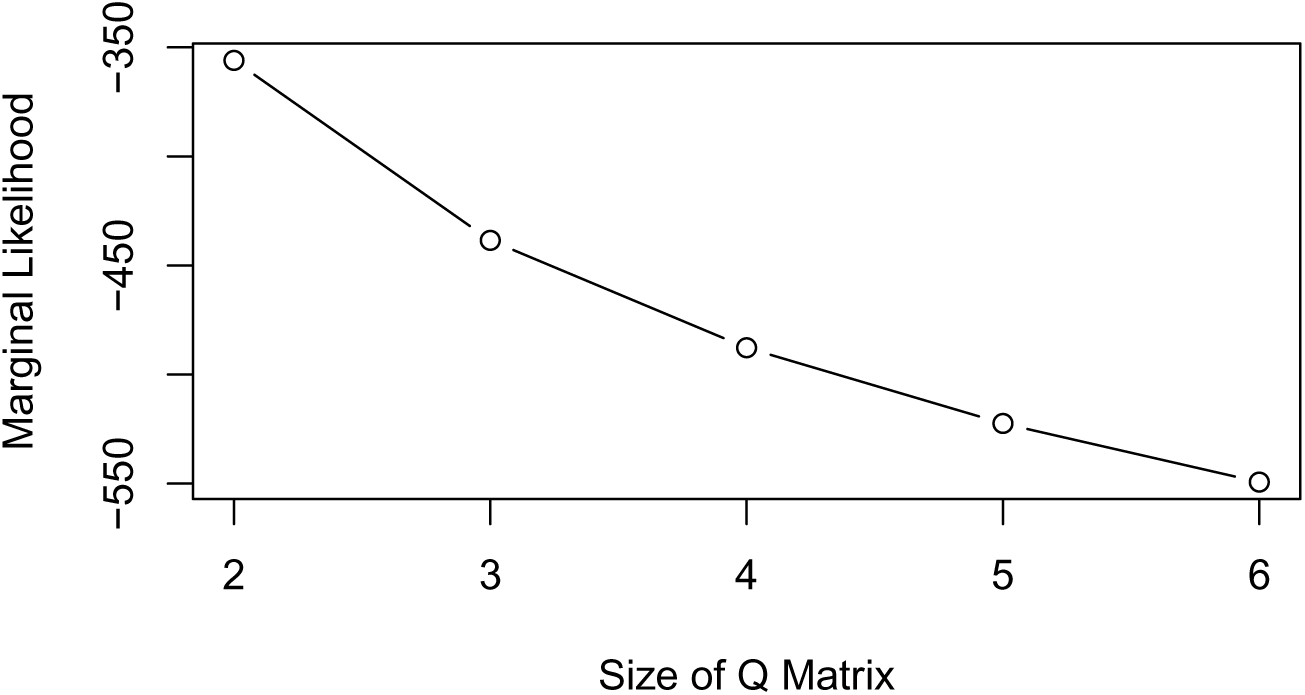
The impact of increasing the size of the Q-matrix on the marginal likelihood calculation. Here a binary alignment was used with stepping stone analysis to calculate the marginal likelihoods. A Q-matrix of size 2-6 was used. As the Q-matrix increased in size, causing binary characters to be in a matrix that was too large, the likelihood decreases.

## Notes

### Competing Interest Statement

The authors have declared no competing interest.

### Summary of Updates

Updated the link to the revbayes tutorial.

